# The penetrant chordoid glioma PRKCA mutation is an oncogenic gain-of-function kinase inactivation eliciting early onset chondrosarcoma in mice

**DOI:** 10.1101/2025.05.28.656646

**Authors:** Véroniqueo Calleja, Jack C. Henry, Mathias Cobbaut, Joanne Sewell, Karine Rizzoti, Francesca Houghton, Stefan Boeing, Nneka Anyanwu, Sunita Varsani-Brown, Thomas Snoeks, Alejandro Suárez-Bonnet, Simon L. Priestnall, Neil Q. McDonald, Angus J. M. Cameron, Peter J. Parker

## Abstract

The penetrant PRKCA D463H mutation, a biomarker and potential driver in chordoid glioma, was found to provoke the development of chondrosarcomas in heterozygous knock-in mice. This mutation entirely abrogates kinase activity, but strikingly no oncogenic phenotype is observed for the related inactivating mutation D463N indicating that the lack of activity is not the driver. In cells, the D463H mutant closely mirrored PKCα WT behaviours and retained ATP binding, contrary to the related D463N mutant. Mechanistically, the PKCα D463H mutant protein was found to display quantitative alterations to the PKCα interactome, enhancing association with epigenetic regulators. This aligned with transcriptomic changes which resembled an augmented PKCα expression program, with enhanced BRD4, Myc and TGFβ signatures. D463H dependent reduced sensitivity to the BET inhibitors JQ1 and AZD5153 indicates the functional importance of these pathways. The data show that D463H is a dominant gain-of-function oncogenic mutant, operating through a non-catalytic allosteric mechanism.

**One Sentence Summary:** A PKCα catalytic inactivating mutation confers gain-of-function properties *-* a paradigm shift in kinase actions.

## INTRODUCTION

There is a breadth of roles assigned to PKC family members in cancer, with examples of both tumour suppressor and tumour promoter actions for individual isoforms (*1–4*). The complexities surrounding this family of kinases in part reflects isoform specificity and context, however underlying discrepancies may also reflect preconceptions. An intriguing example of this is the role of PKCα in chordoid glioma. In this circumscribed, low-grade but difficult to access CNS tumour, two groups published the finding of completely penetrant heterozygous clonal mutation in *PRKCA* (*5, 6*). The clonal somatic mutation repeatedly identified was a transversion (c.1387G>C) that alters the catalytic aspartate of PKCα to a histidine residue (D463H). However, no LOH in the 17q24.2 locus, nor other consistent mutations, deletions, amplifications associated with *PRKCA* and no mutations in other oncogenes or tumour suppressors were reported for these tumours. The same D463H mutation was also documented by other groups in a 16 patient clinical study (*7*) and in two case studies (*8, 9*), suggesting the mutation is a specific diagnostic marker of chordoid glioma. Intriguingly, extrapolating from the pioneering work on PKA (*10, 11*) this *PRKCA* mutation would be an inactivating mutation, hence the parsimonious conclusion would be a loss-of-function and a suppressor role for *PRKCA.* However, the heterozygous occurrence of the D463H mutation in chordoid glioma does not align with a classical suppressor role unless through dominant loss-of-function or haploinsufficiency. Such assumptions would seem nevertheless provocative as there is only one mutation found (D463H mutation) while there are many ways to inactivate PKCα.

Understanding the ‘sense’ of action of this penetrant *PRKCA* mutation and the apparent lack of other driver mutations, would offer vulnerability and opportunity to a non/post-surgical intervention in chordoid glioma. To address this, we used genetic models and comparative biochemistry. Here we show that a global heterozygous knock-in of the D463H mutation of *Prkca* in mice drives chondrosarcoma formation in very young neonates. We determined that the *Prkca* D463H mutation is a non-catalytic gain-of-function oncogenic mutation which drives a transcriptomic program resembling an augmented WT response. However, the catalytic activity of PKCα *per se* is not the trigger of the phenotype as a related knock-in inactivating mutation of the same residue (D463N) has no such effect either in a heterozygous or homozygous setting. The distinct behaviour of the D463H mutant is reflected by mutant specific alterations to the PKCα interactome, linking to transcriptomic changes and demonstration of a functional link to BRD4. A model of action of this unique PKCα oncogenic mutant is presented, impacting our wider appreciation of kinase behaviour, and prescribing a distinctive approach to intervention.

## RESULTS

### The heterozygous expression of *Prkca* D463H mutant in mice has a dominant harmful effect causing hindlimb deformation and neonatal death

To assess the in vivo impact of the PKCα D463H mutation we knocked-in a single allele into C57BL/6J mice using two different guide RNAs (Fig. 1A). From 3 litters (table S1), 8 pups harboured the mutation (from 11% to 90% penetrance) of which all but 3 had to be culled within 2 weeks due to ill health. Only one male (22% penetrance) survived to breeding age. The WT mice from the same cohorts were healthy, suggesting that the presence of the *Prkca* mutation rather than a side effect of the CRISPR/Cas9 directed knock-in, was responsible for the morbidity of the animals.

**Fig. 1.**
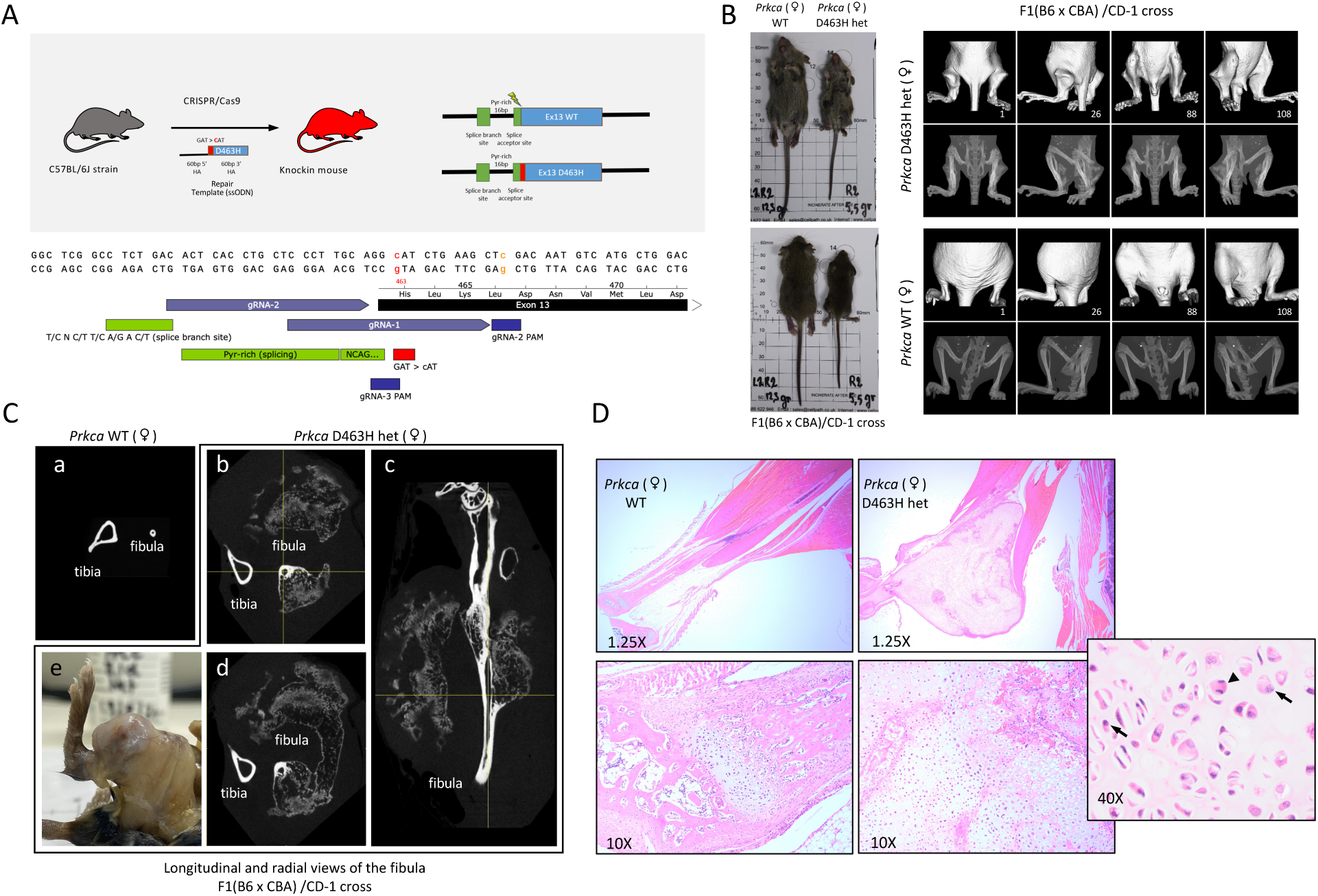
heterozygous knockin mice for PKCα D463H mutation develop chondrosarcoma. (**A**) CRISPR/Cas9 strategy to introduce the D463H point mutation in C57BL/6J mice strain. (**B**) Photos, and scan of representative WT and D463H het mutant mice in ventral and dorsal positions showing the extensive loss of weight and small size of the D463H het mutant compared to the WT. The CT-scans focus on the hindlimbs and present multiple views of the bilateral leg deformation of the D463H het mutant mouse (top panels) compared to the WT (bottom panels). (**C**) Photo of the leg (e) and high-resolution CT-scans showing the aberrant growth of cartilage around the fibula of the D463H mutant mice (transversal view for two different D463H het mice (b and d) as well as an (orthogonal view (c)) and not on the WT (panel a). (**D**) H&E stained sections of WT and D463H het mouse hindlimbs. A disorganized, histologically malignant, chondrocyte cell proliferation characterizes the abnormal phenotype in the D463H mutant mice. In the 40x image the arrowhead is pointing to a mitotic figure and the arrows to cells showing cellular atypia like anisokaryosis and anisocytosis.

In timed matings of the one surviving male mosaic mouse with C57BL/6J females (table S2), 7 heterozygous mice were identified out of 46 pups from 6 different litters, indicating germline transmission and a roughly Mendelian segregation given the mosaic nature of this founder. The *Prkca* D463H heterozygotes pups were smaller or hunchbacked and were malnourished, needing to be culled at about 3 weeks despite the addition of high energy mash nutrition to encourage weight gain.

To determine whether the C57BL/6J mouse background might have contributed to the phenotype, we used outbred CD-1 mice that carry greater genetic diversity, are a more robust strain and produce larger litter sizes than inbred C57BL/6J mice (table S3). Using sperm from the surviving male mosaic mouse, 3 heterozygotes from 4 litters were born. Despite the genetic background change the pups again showed signs of sickness with hindlimb deformation, typically becoming symptomatic at about 2 weeks of age. However, one het male survived longer (11 weeks) and was crossed with CD-1 females producing 4 heterozygotes in 12 progenies (table S4), none of which survived to breeding age, but all of which displayed malnourishment and a hindlimb phenotype (see further below).

The intrinsic constraint on breeding and back-crossing to link genotype to phenotype, led us to perform a second round of gene editing. New mosaic mice were obtained from 8 litters on a C57BL/6J background (4 with guide RNA1 and 4 with guide RNA2) and 8 litters on an F1(B6 x CBA) background (4 with guide RNA1 and 4 with guide RNA2). From these litters 56/103 mice born were mosaic and as observed in the first round of CRISPR knock-ins the mosaic mice but not their WT littermates were malnourished and displayed deformation of the hindlimb. One mosaic male mouse from C57BL/6J background and one male from the F1(B6 x CBA) background that expressed 12% and 18% mutation penetrance respectively were selected for breeding (table S5). Both mice were crossed with two WT CD-1 females producing 15/39 mice from the C57BL/6J male and 6/23 mice from the F1(B6 x CBA) male heterozygous for the D463H mutation. Yet again from the 5 litters obtained (62 mice in total) all the 21 heterozygotes were sickly and displayed a deformation of the hindlimbs. These findings support the conclusion that the heterozygous mutation was a trigger for the malnourished, hindlimb phenotype observed.

### The *Prkca* D463H mutation elicits bilateral hindlimb chondrosarcoma

To address the penetrant hindlimb phenotype of the D463H heterozygotes, representative animals were imaged by CT (Fig. 1). While the CT-scan of the hindlimb showed the extensive loss of weight and the bilateral leg deformation (Fig. 1B), no deformation was visible on the forelimbs (fig. S1A). Furthermore, CT scan of skulls showed a small decrease in size for the heterozygous mice which was not significant when related to overall body size and weight (fig. S1B).

The longitudinal and radial cross sections of the hindlimbs of representative D463H heterozygotes when compared to a WT littermate clearly showed the aberrant growth of cartilage and bones (Fig. 1C). Effacing the gracilis, semimembranosus, semitendinosus, and gastrocnemius muscles there were bilateral, symmetrical, mesenchymal neoplasms, composed of moderately differentiated chondroblasts and chondrocytes surrounded by a pale cartilaginous matrix. There was moderate anisokaryosis and anisocytosis and only a single mitosis was observed in ten high-power fields (400x). The bulk of these neoplasms blended with an outer rim of poorly mineralized woven bone with a population of activated osteoblasts. Occasionally within the neoplasms there were entrapped segments of nerve fibres. There was no evidence of chronic inflammatory changes or haemorrhage. Forelimbs and the remaining soft tissues were histologically unremarkable. Based on both the histological appearance and the infiltrative nature of these proliferative lesions, they were consistent with well-differentiated chondrosarcomas (Fig. 1D). These lesions were associated with morbidity due to muscle and nerve compression and atrophy resulting in the disuse of the hindlimbs. The loss of weight was attributed to the lack of easy access to food due to the competition with the other pups for the mother’s milk. Furthermore, sections from both forelimbs and hindlimbs, lung, heart, liver, kidney, and spleen of WT littermates were also examined and were histologically unremarkable.

Since the PKCα D463H mutation had initially been identified in chordoid glioma tumours originating from β2 tanycytes in the third ventricular region (*5, 6*) (fig. S1C), we analysed the brains of D463H mutant mice and wild-type controls at the level of the median eminence. As the D463H mutation did not allow the survival of the het mice beyond a few weeks of age, the analysis of the adult brain could only be performed on a single mosaic mouse (IRCP 4.1e). This mosaic female (30 weeks - 23% D463H penetrance) was the only surviving littermate of the mosaic founder male IRCP 4.1c (table S1). Additionally, 2 week old D463H heterozygotes and wild-type control mice were analysed (fig. S1D, E). These analyses did not detect any change in the brain architecture or aberrant growth in any of these animals.

### PKCα D463H mutant is inactive

In order to determine the mechanism by which PKCα D463H mutant was driving tumour formation, we first analysed the impact of the D463>H change on kinase activity. Recombinant WT and mutant full-length proteins (Fig. 2A and fig. S2) and isolated kinase domains (Fig. 2B) were assessed for protein kinase catalytic activity with both peptide and protein substrates. As a comparison, the behaviour of a previously characterised mutant at the same site (D463N) was analysed; this mutant had previously shown lack of catalytic activity but retention of protein integrity (*12*). Characteristic activity was observed for the WT protein, however neither D463 mutant displayed any phosphotransferase activity (Fig. 2C-E). Assessing ATPase activity showed that this was detectable for the WT protein, albeit orders of magnitude lower than phosphotransferase activity, but again, no activity was observed for either D463 mutant (Fig. 2D). To ensure that there was no compensating function for PKCα derived from mammalian cell expression, we also determined the activities of these proteins on expression in HCT116 (colorectal cancer derived) and U87MG (malignant glioma derived) cells. Using PKCα pull-down/kinase assays a series of phosphorylated coprecipitated proteins could be detected, but only for WT and not for either of the D463 mutant proteins (Fig. 2F and 2G).

**Fig. 2.**
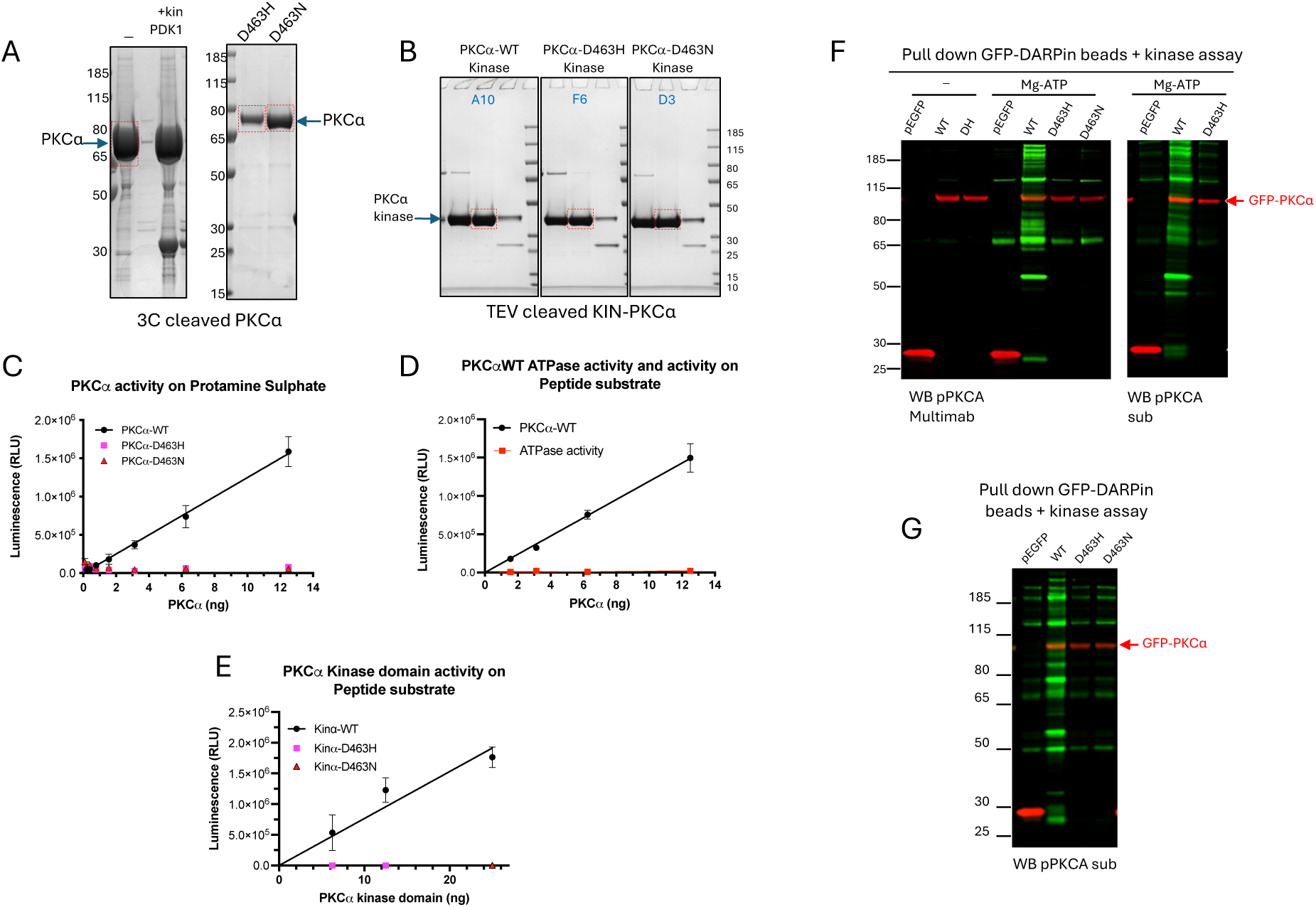
Full length and kinase domain PKCα D463H mutants are kinase dead. (**A**) expression of recombinant full-length WT-PKCα and mutants D463H and D463N after C3 cleavage. The co-expression of PDK1 kinase domain together with WT-PKCα in insect cells is not necessary to stabilise further the expression of the protein. (**B**) expression of recombinant WT-PKCα and mutants D463H and D463N kinase domains after TEV cleavage of full-length recombinant PKCα. The fraction used for the experiments is highlighted in red. Kinase activities were measured using the ADP-Glo method (C-F). (**C**) Recombinant full-length PKCα activity in vitro on protamine sulphate. The kinase activity of WT is of 523 nmol/min/mg. No activity is detected for both D463 mutants (**D**) Recombinant full-length PKCα activity measured in vitro on a peptide substrate (257 nmol/min/mg on PPSS). There is no unspecific ATPase activity unless at very high saturating PKCα concentration. (**E**) PKCα recombinant isolated kinase domain activity in vitro measured on a peptide substrate. The kinase activity of WT kinase domain is of about 300 nmol/min/mg. No activity is measured for both D463 mutants. (**F**) Co-precipitation of associated PKCα substrates using DARPin pull down of GFP-tagged PKCα WT and mutants expressed in U87MG cells. The substrates ladder is detected by western blot using two different anti phospho-PKCα substrate specific antibodies (pPKCA sub and pPKCA multimab in green). The expression of PKCα is visible (red) using an anti GFP antibody. The phosphorylation of the associated substrates is only detected upon in vitro kinase assay using Mg-ATP, calcium, and lipids activators (phosphatidyl serine). The mutants of the residue D463 do not have kinase activity and do not show specific substrates’ bands phosphorylation. (**G**) Co-precipitation of associated PKCα substrates using DARPin pull down of GFP-tagged WT-PKCα and mutants expressed in U87MG cells. Unlike WT, no substrates’ phosphorylation (WB pPKCA sub) can be detected with the mutants D463H and D463N upon kinase assay.

### *Prkca* D463N mutation does not phenocopy *Prkca* D463H in mice

Notwithstanding the lack of an overt tumour driver phenotype in hetero- or homozygous knock-out *Prkca* mice (*13*), in principle the effect of the D463H inactivation mutation might still be compatible with a dominant negative impact on a suppressor *Prkca* allele as opposed to a gain-of-function allele. To address this question, we knocked in the *Prkca* D463N inactivating mutation not found in chordoid gliomas but known to retain protein integrity (*12*). We used the same guide RNAs as described for the D463H mutation but with a modified repair template to change codon D463 to an asparagine (see Materials and Methods). From the 16 C57BL/6J mosaic mice obtained with the D463N mutation (table S6), 3 were bred to assess germline transmission, yielding 15 of 30 pups that were *Prkca* D463N heterozygous from 4 litters (table S7). All mice were asymptomatic and 8 of these offspring were crossed with C57BL/6J, F1(B6 x CBA) or CD-1 mice. In these different backgrounds, a Mendelian ratio of 30 heterozygotes out of 58 pups were obtained, and all the 30 heterozygous *Prkca* D463N mice obtained from all backgrounds survived for at least 20 weeks without any health issues (table S8). Hence the monoallelic expression of kinase inactive *Prkca* D463N is not sufficient to drive the chondrosarcoma phenotype.

It remained possible that the D463N mutation might be haploinsufficient failing to ‘dominate’ the WT allele in the manner that the D463H mutation might be argued to do. To address this, we bred the D463N allele (C57BL6 background) to homozygosity (table S9). 12 homozygotes, 14 heterozygotes and 6 wild types were generated. None of the mice displayed any phenotype and the histology of organs was entirely normal. It is noted that neither mutant appears to express well in mouse tissues with the heterozygote D463H+WT expressing 38% normal PKCα protein (range 15-65) and the D463N+WT expressing 48% (range 41-53); we cannot distinguish WT and mutant proteins so cannot define contributions to assess if D463N versus D463H levels impact phenotype. However distinctions from ex vivo studies (see below) demonstrate distinctive biological properties.

The rapid onset of chondrosarcoma in the *Prkca* D463H mutant heterozygotes but not in the *Prkca* D463N homo- or heterozygotes indicates that the D463H mutation is a positive driver of neoplastic disease. This suggests that the mutant either has neomorphic properties or that it is a gain-of-function PKCα allele that redefines effector outputs of this protein kinase.

### PKCα D463H behaviour aligns with WT-PKCα properties

With a view to understanding how the D463H mutant varies from the D463N mutant and whether this tracks with WT behaviour or represents neomorphic behaviour, we analysed the effects of the D463H and D463N mutations on biochemical and in cell properties. Thermal shift assays and microscale thermophoresis were conducted to assess whether differences between the mutants might reflect distinct nucleotide binding properties. The mutant proteins showed monophasic melting curves, in thermal shift assay, unlike WT full length protein which displayed some heterogeneity in its behaviour (Tm WT1 and WT2). However, all proteins displayed a shift in melting temperature(s) on binding ATP or ADP (fig. S3A) with some variations in the extent of nucleotide stabilisation (fig. S3B). Despite the biphasic melting behaviour of the WT protein complicating comparisons with the mutants, there is a substantial reduction in ATP affinity for the D463N mutant (shown by higher EC50) compared to the WT with the D463H mutant displaying a more modest loss of affinity. Both mutants displayed a similarly reduced affinity for ADP relative to the WT protein (fig. S3C). To address this differential loss of affinity for ATP more directly, we employed an orthogonal approach, assessing ATP binding to full-length recombinant PKCα and mutants in vitro with microscale thermophoresis (MST). This revealed that both D463H and D463N bind ATP less well than the WT (Kd^WT^= 0.70 ± 0.23 µM) (fig. S3D); critically the D463N bound ATP with very much lower affinity (Kd^D463N^=1.5±0.7mM) than D463H (Kd^D463H^=94±17µM) μM. These values predict significantly greater ATP occupation of the D463H mutant compared with the D463N mutant at physiological ATP concentrations.

To compare cellular behaviours of the PKCα WT and mutants, GFP-fusion proteins were expressed in U87MG cells, and characteristic activation-induced degradation monitored (Fig. 3A, B). All three GFP-tagged proteins were phosphorylated at their priming sites (Fig. 3A panel b) indicative of protein integrity, consistent with previously published findings for the D463N mutation (*12*). There was however a modest decrease in the phosphorylation of both mutants on the activation loop (about 25%) compared to WT (Fig. 3A panels a and b) perhaps reflecting a more open conformation (see below). The dephosphorylation and degradation of PKCα known to occur upon prolonged phorbol ester-dependent activation (Fig. 3B) was observed for WT and D463 mutants. Notably, the kinetics of the D463H mutant response mirrored that of PKCα WT whilst the D463N mutant was slower both in the dephosphorylation of the HM motif (Fig. 3A panel c), turn-motif and the activation loop (fig. S4A, B). The D463H mutant matched with WT in its degradation (fig. S4E). Myc-fusion proteins similarly showed the D463H mutant following the WT kinetics of induced dephosphorylation and degradation with the D463N mutant being delayed (Fig. 3C and fig. S4C, D).

**Fig. 3.**
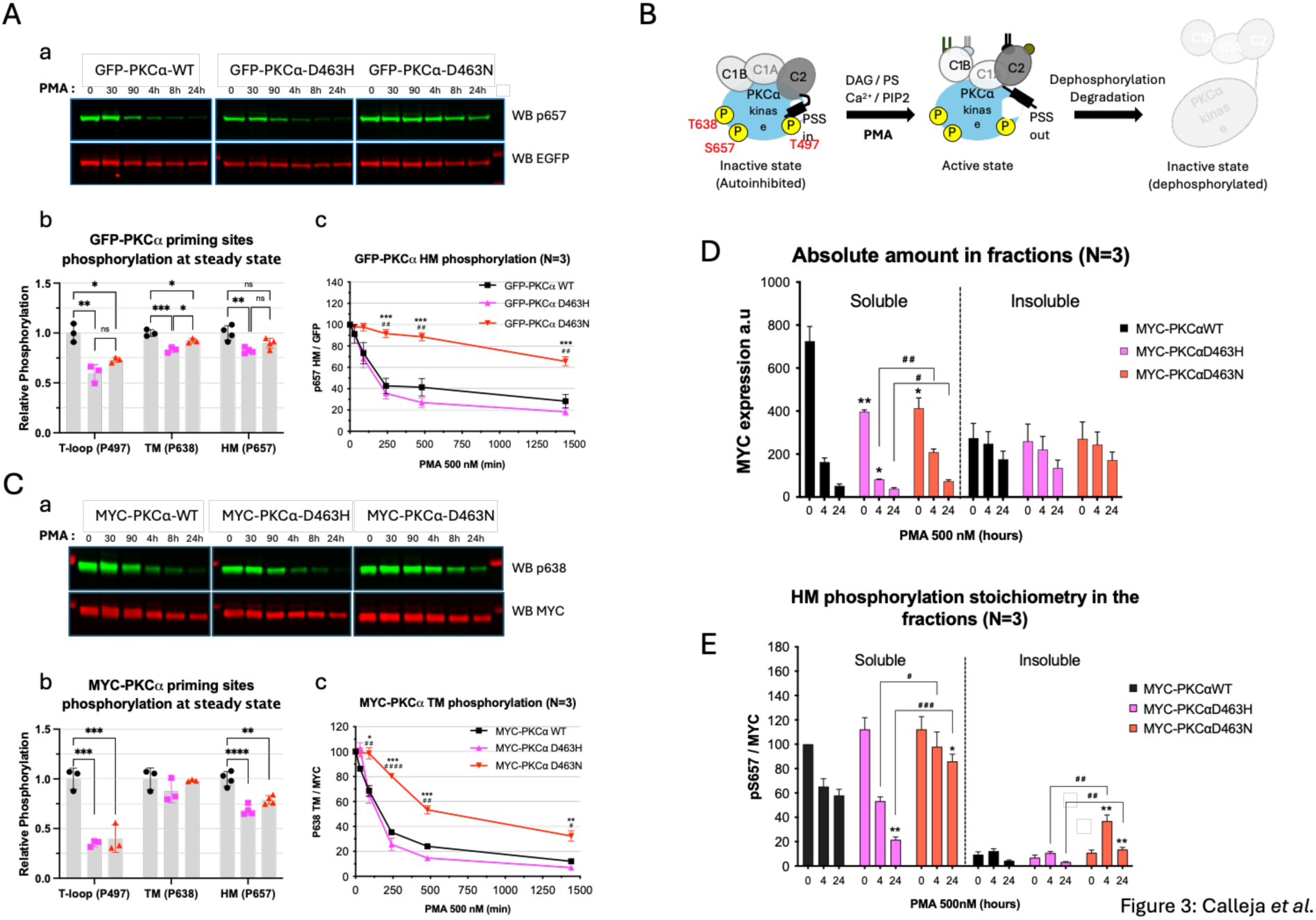
PKCa D463H mutant follows WT-PKCα regulation. (**A**) U87MG cells expressing transiently GFP-PKCα wild-type and mutants are treated with PMA up to 24 h. Panel (a) the time course decrease in the phosphorylation of the HM motif (pS657) and GFP-PKCα expression upon PMA is detected by western blot with a phospho-specific antibody (green) and an anti-GFP antibody (red). Panel (b) shows the steady state phosphorylation of the three PKCα WT and mutants activation sites (T-loop, TM and HM) prior to PMA stimulation. A one way Anova was used to calculate the significance of the variation between WT and mutants(n=3). Panel (c) shows the decrease in HM phosphorylation upon PMA treatment and an unpaired t-test was used to compare the HM phosphorylation of D463N vs D463H (p-values*) or WT (p-values^#^) (n=3). The data in (b) and (c) are expressed as a percentage of decrease relative to the WT signal. (**B**)Schematic representation of the life cycle of PKCα upon activation or PMA treatment. PKCα is in an autoinhibited inactive state but in a fully phosphorylated (primed) conformation prior to stimulation. Upon activation the protein translocates to the plasma membrane where the regulatory domains (C1A/C1B and C2) bind to the lipids where PKCα becomes fully active when the pseudo substrate (PSS) is ejected from the catalytic site. The open conformation becomes substrate for the phosphatase that dephosphorylate the priming sites and renders PKCα prone to degradation. (**C**) U87MG cells expressing transiently Myc-PKCα wild-type and mutants are treated with PMA up to 24 h. Panel (a) The time course decrease in the phosphorylation of the turn motif (pT638 TM) and expression of Myc-PKCα upon PMA is detected by western blot with a phospho-specific and anti-Myc antibody. Panel (b) shows the steady state phosphorylation of the three activation sites (T-loop, TM) and HM) of WT-PKCα and mutants prior to PMA. A one way Anova was used to calculate the significance of the variation between WT and D463H or D463N (n=3). Panel (c) shows the decrease in TM phosphorylation upon PMA treatment (n=3). An unpaired t-test was used to compare the TM phosphorylation of D463N vs D463H (p-values*) or WT (p-values^#^). The data in panels (b) and (c) are expressed as a percentage of decrease relative to the WT signal. (**D**) Triton-X100 fractionation of U87MG cells expressing Myc-tagged WT-PKCα and mutants following PMA treatment for 4h or 24h (n=3). The soluble and insoluble fractions were analysed by western blot using an anti Myc antibody to calculate the absolute amount of protein upon degradation. An unpaired t-test was used to calculate the significance of the changes between WT and D463H or D463N (p-values*) or D463H vs D463N (p-values^#^). (**E**) Soluble and insoluble fractions were also analysed by western blot using an anti pS657 (HM) and the stoichiometry of HM phosphorylation was calculated for each PKCα and mutants. An unpaired t-test was used to calculate the significance of the changes in the soluble and insoluble fractions. In all the experiments the significance was assessed using a t-test P<0.05 (*), P<0.01 (**), P<0.001(***), P<0.0001 (****).

The distinction between the D463 mutants was also reflected in the detergent soluble versus insoluble distribution of proteins (Fig. 3D), where the D463N mutant was retained in the neutral detergent soluble fraction and was dephosphorylated more slowly in this context (Fig. 3E); in contrast the D463H mutant behaviour closely aligned with the WT protein.

### Evidence for distinct protein interactions for PKCα D463H

The distinctive in vivo behaviour of D463H with respect to WT and D463N, led us to assess how this specific mutant might impose itself to trigger oncogenic signals in a non-catalytic fashion. To address this, we first sought to examine differences in the interactome between WT and D463 mutant PKCα in U87MG cells. U87MG cells were transfected with Myc-tagged WT-PKCα, PKCα-D463H, PKCα-D463N or empty-vector control as technical triplicates; all PKCα constructs expressed to comparable levels and were recovered equally in Myc immunoprecipitates (Fig. 4A), which were subsequently analysed by tandem MS/MS. Any proteins recovered in precipitates from empty-vector transfected cells were excluded as artefacts. A total of 200 proteins were identified as potential PKCα interactors, including known PKCα interacting proteins (Caveolin (*14*); PKCγ (*15*); Annexin V (*16*)). Principal component analysis and unsupervised clustering of interactomes indicates a clear distinction between the interactomes of WT, D463H and D463N mutants, with the D463H mutant interactome showing greater similarity to WT-PKCα (Fig. 4B and 4C) than D463N. Approximately 90 proteins were identified as interactors with both WT and PKCα-D463H, with 15 showing statistically significant (P<0.05) differences in their abundance between replicates (Fig.4D and table S12). Further, 11 high confidence interactors were exclusively recovered with PKCα-D463H, including the oncogenic BET domain containing epigenetic regulators BRD3:BRD4 (table S13); BRD4 has been implicated previously as a driver of glioma progression, potentially through promotion of Myc driven anabolic pathways (see (*17, 18*)). To further explore PKCα functional roles for PKCα or the PKCα-D463H mutant in glioblastoma biology, STRING (*19*) analysis was conducted on proteins showing preferential binding to D463H over WT, which revealed networking between epigenetic regulators, including BRD4 and the MLL1 complex components HCFC1 (HCF-1) and PRPF31 (Fig. 4E). These pathways are associated with promotion of RNA polymerase II and Myc driven transcription, through permissive histone modifications (*20–25*). These distinctive binding patterns suggest a gain-of-function for the D463H mutant and importantly, show clear differences between the D463H and D463N mutants.

**Fig. 4.**
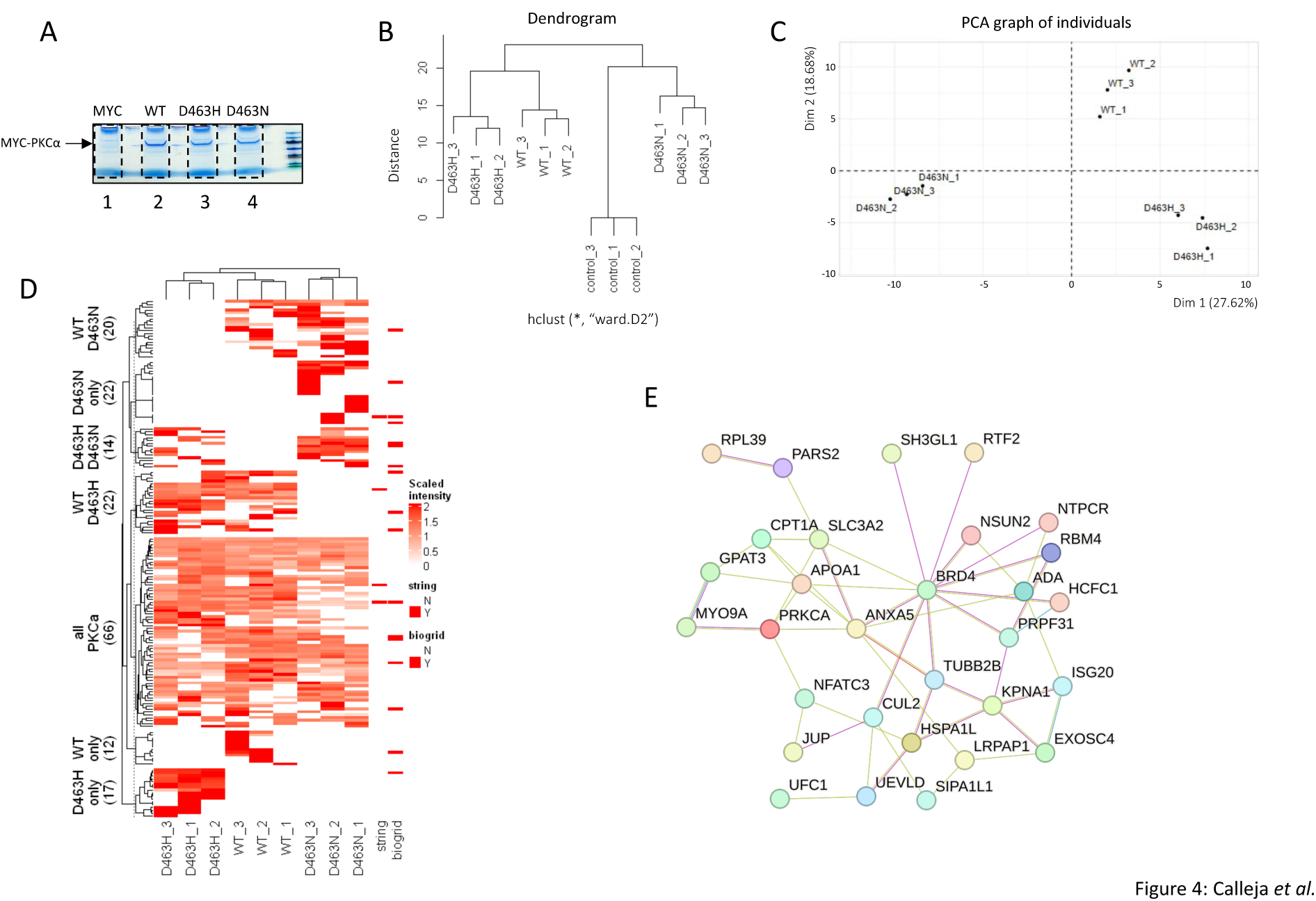
Proteomics of WT and mutant PKCα pull downs. (**A**) WT and mutant proteins as indicated were expressed as Myc-fusions and captured on anti-MYC beads (described in Materials and Methods). Proteomic analysis of captured proteins was performed by tryptic fragmentation and mass spectrometry and two-way comparisons were made. (**B**) Dendrogram… (**C**) Principal component analysis of the interactors found in each PKCα WT and mutants showing the spatial separation between the Myc-PKC WT and mutants and between the two mutants. STRING analysis on proteins preferentially recovered in D463H immunoprecipitates compared with WT-PKCa immunoprecipitates. (**D**) Heatmap presenting the mutants only interactors (11/17 high confidence hits in D463H only interactors) or interactors found all PKCα or in pair combinations of WT and PKCα mutants. (**E**) STRING network is displayed at 0.15 confidence level demonstrating networking of key epigenetic regulators, including BRD4 and the MLL1 complex components HCFC1 and PRPF31, and their relationship to PKCa (PRKCA). Known interactions from experimentally determined (magenta) or curated datasets (cyan) are indicated, along with associations from text mining (green).

We employed TurboID-PKCα fusions (fig. S5A, B) in U87MG cells with or without PMA activation to identify by western blot biotinylated partner patterns using fluorescently labelled streptavidin. The fusion proteins themselves were biotinylated as expected (fig. S5C) alongside a significant number of bands identified by the streptavidin probe (fig. S5D). This technique allowed us to detect a prominent biotinylated species of about 27 kDa in the D463H expressing cells that was absent from the WT cells under basal conditions but became biotinylated on stimulation with the PKCα activator PMA (fig. S5D white star). This is indicative of a specific up-regulated interaction for the D463H mutant at steady state, which mimics the up-regulated interaction observed on acute allosteric activation of the WT protein. Both the steady state interactome studies and the TurboID proximity analyses support the concept that D436H alters interaction of PKCα with binding partners to drive an augmented and functionally distinct oncogenic program. Enhanced association with epigenetic regulators also helps explain how a single point mutation might induce profound oncogenic reprogramming.

Structure predictions provided insight into the molecular details underlying this privileged behaviour. In the case of PKCα, AlphaFold3 yielded confident predictions for the effect of a single-residue Asp-463 substitution. We predicted the structure of WT, D463H and D463N mutants, generating 100 models for each protein. For the WT, the top scoring model displayed an auto-inhibited conformation, with the pseudosubstrate sequence reliably positioned docking into the active site in a substrate-binding mode (see PAE matrix fig. S6A), and the C1a contacting αB, the αB-αC loop as well as the C-tail (fig. S6A). The top scoring model for D463N and D463H however displayed a confident prediction of a pseudosubstrate-out conformation with the C2 domain packing against the back of the C-lobe of the kinase domain and the C-tail, and the C1 domains positioned under the C-lobe. Mg^2+^ ions (one in D463N, both in D463H) were modelled out of the active site in the C2 domain, and the kinase domain was in a more open conformation, while still more closed compared to apo predictions. We then broadened the analysis to all predicted models. Inspection showed them all to be either in PS-in or PS-out state, so we first classified them based on their predicted conformation. For the WT protein, most models (88%) were predicted as PS-in, whereas most were predicted as PS-out for D463H (96%) and D463N (81%) (fig S6B). We next compared model quality and characteristics of the predicted models by comparing the pTM score and median PAE score. The latter is used as a metric to compare the confidence in the relative positioning of domains within a predicted conformation. The mutant kinases showed better pTM scores in the PS-out conformation (fig. S6C) and a better confidence in this conformation (fig. S6D), with the best models observed for the D463H mutant. In the PS-in conformation however pTM scores were better for WT. The D463N mutant showed a bimodal distribution, with about half of its PS-in models predicted with WT or D463H-like scores (fig. S6E). Similarly, the confidence in PS-in conformers was worse for the mutants (fig. S6F). Overall, the WT protein is predicted as a PS-in conformer with high confidence, whereas the D463H mutant is predicted as a PS-out conformer with high confidence. These predictions suggest the D463H mutant presents as a PS-out, disinhibited state that is effector-competent, similar to an activated form of the WT protein, albeit incapable of ATP hydrolysis. The D463N mutant shows similar characteristics as the D463H mutant, although less pronounced, and its functional characterization supports a different nucleotide-binding profile that may not be compatible with effector trapping. The effect of D463 substitution is also supported by a predicted effect on stability of the WT kinase which is more pronounced for D463H (ΔΔG^WT>D463H^ -2.1 kcal/mol, ΔΔG^WT>D463N^ -1.4 kcal/mol).

### PKC**α** D463H is a gain-of-function PKC**α** allele

Having established distinct biochemical characteristics and protein interactomes for the oncogenic PKCα D463H mutant, we next examined transcriptional changes in the U87MG models to explore mechanisms driving transformation. We compared U87MG glioma cells stably expressing WT-PKCα or the two D463 mutants. mRNA expression of PKCα was comparable between WT and the two mutants (Fig. 5A), however, the expression of D463N protein was reduced compared to WT or D463H (Fig. 5B). This may reflect differences in the protein stability or proteasomal targeting for the distinct D463 mutants. Nevertheless, all expressed proteins altered the pattern of mRNA expression and principal component analysis indicated tight clustering for biological replicates for each construct and the parental samples (Fig. 5C). Furthermore, all constructs resulted in significant displacement in principal component space away from the parental samples indicating that despite the difference in expression levels (in D463N versus the other two) there is a distinct impact of each PKCα allele on mRNA profiles.

**Fig. 5.**
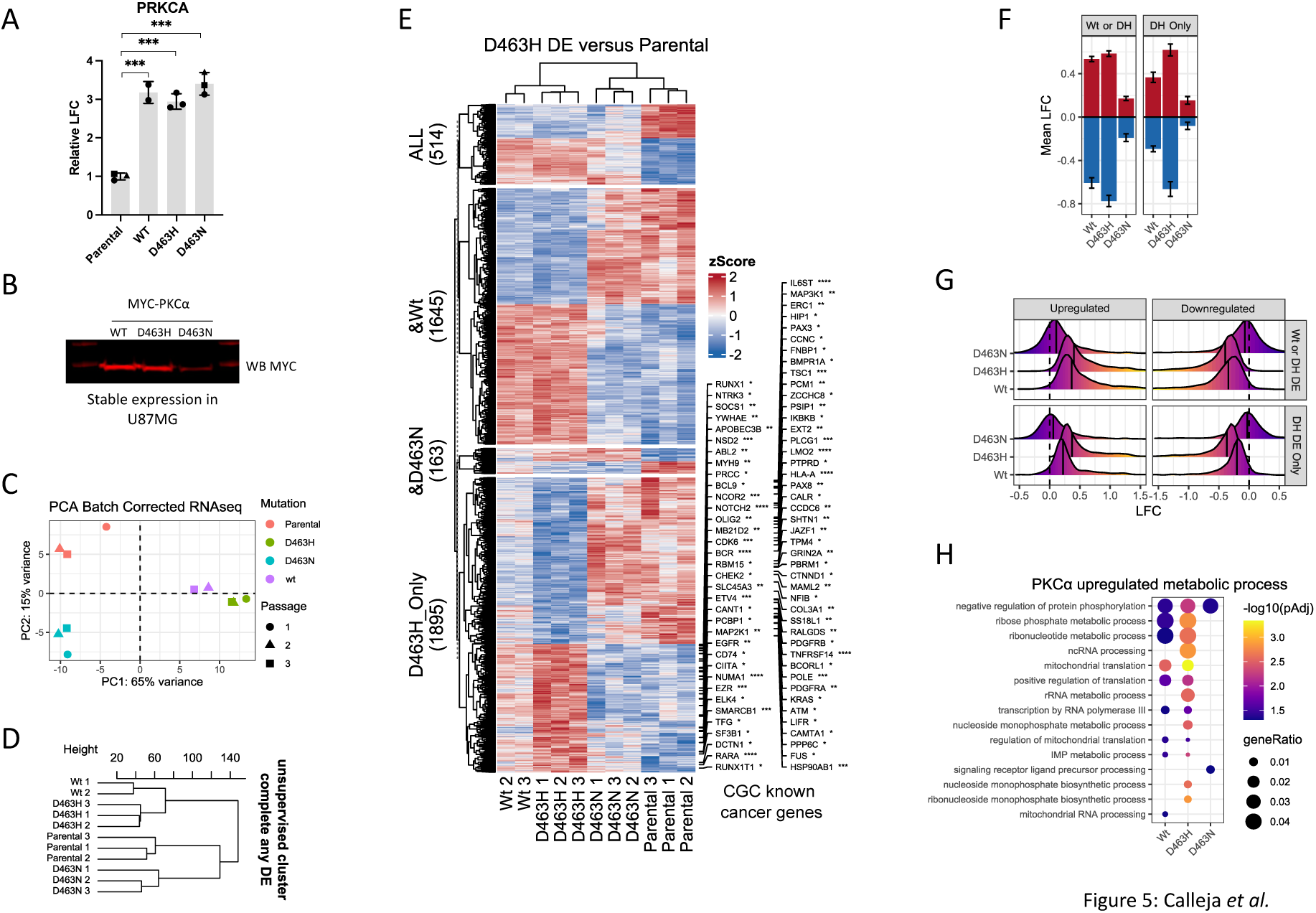
PKCα D463H is a dominant gain-of-function mutant which drives an augmented WT-PKCα transcriptomic signature. (**A**) mRNA expression of PRKCA in stable U87MG cells with Myc-tagged WT-PKCα, D463H and D463N mutants determined by RNA sequencing. Statistical comparison to parental cells was performed using an unpaired t-test P<0.001 (***) (n=3). (**B**) Stable protein expression of Myc-tagged WT-PKCα and mutants D463H and D463N in stable U87MG by western blot using an anti Myc antibody. (**C**) Principal component analysis of batch-corrected mRNA expression from the different stable cell lines expressing Myc-tagged WT-PKCα and mutants. (**D**) Unsupervised hierarchical clustering analysis of gene expression from all significantly differentially expressed genes in stable U87MG cells with WT-PKCα, D463H and D463N mutants when compared with parental cells. (**E**) Heatmap of genes differentially expressed with overexpression of Myc-PKCα D463H mutant when compared with parental cells. Tiles are coloured by per-gene z-scores across all samples. Rows are clustered by overlap in significant differential gene expression with WT and D463N samples when compared to parental cell lines. Cancer gene consensus (CGC) known cancer genes are annotated for the D463H differentially expressed only cluster. (**F**) and (**G**) Mean log fold change (F) and distribution of log fold change (G) for the WT-PKCα, D463H and D463N mutants when compared with parental cells for genes significantly differentially expressed in WT or D463H mutant and D463H mutant only when compared with parental cells. (**H**) Overrepresentation analysis of the top eight significant gene ontologies from the metabolic process (GO:0008152) gene ontology category child terms for WT-PKCα, D463H and D463N mutant samples. Upregulated differentially expressed genes in stable U87MG cells with WT-PKCα, D463H and D463N mutants when compared with parental cells were used for overrepresentation analysis.

Remarkably, the D463H mutant RNAseq data segregated closely with the WT and well away from the D463N mutant (Fig. 5C), consistent with similarities in biochemical behaviour and interactome changes. Unsupervised hierarchical clustering of gene expression profiles also indicates the close relationship of WT and the D463H mutant (Fig. 5D), which entirely mirrors that seen for the proteomic analysis (Fig. 4B). While the majority of differentially expressed (DE) genes induced or suppressed by WT-PKCα follow the same directionality for PKCα D463H, the amplitude of change is generally greater for the D463H mutant; this can be seen in both principal component analysis, scatter analysis and through clustering of all DE genes when compared with parental (Fig. S7A, B). Exploring the amplitude differential between WT and the D463H samples, we separated gene clusters according to their significant DE expression patterns (relative to parental) between the WT, D463H and D463N experimental conditions (Fig. 5E). The largest individual cluster comprises 1895 genes which are significantly up or down regulated only by the D463H mutant; within this cluster, the vast majority (94.6%) of genes displace from the parental in the same direction for both the D463H and WT-PKCα conditions, but the mean Log Fold Change (LFC) for D463H is approximately doubled on average for the whole 1895 gene group (Fig. 5F). Examining the distribution of LFC clearly indicates the directional similarity but augmented response of the D463H compared with WT, in contrast to the distinct behaviour of the D463N mutant (Fig. 5G). Numerous known cancer-genes (cancer gene consensus - COSMIC) are present within the D463H only group that have augmented gene expression (Fig. 5G). The second largest cluster comprised genes which are DE for both WT and D463H, and these largely show parallel behaviour, but strong differential expression compared with D463N and parental (Fig. 5E). These dominant groups underly the directionality and amplitude of movement observed in the principal component analysis and also identify a specific gene-expression output which is noticeably augmented by the D463H mutant. To assess functional impact, we conducted geneset over-representation analysis, focussed on genesets associated with translation and metabolism (Fig. 5H). This demonstrates D463H acts as a gain-of-function mutant, augmenting expression of multiple anabolic genesets compared to WT and elicits an entirely distinct pattern compared to D463N.

Only a minor group of 163 genes showed some degree of similarity between the two kinase-inactive D463N and D463H mutants compared with WT (Fig. 5E). We would interpret these alterations as those where PKCα kinase catalytic activity plays an active effector role. Overall, however, these data suggests that PKCα kinase activity is of minor importance to ‘positive’ effector outputs in the context of this glioblastoma derived cell line, consistent with the in vivo observations.

Among <u>the D463H</u> DE genes (Fig. 6A, fig. S7) there are many known and candidate cancer genes, including enhanced expression of NRAS, KMT2D and CD74 (Cancer Gene Census - COSMIC). The majority of these genes are not DE between WT and the D463N mutant. Geneset enrichment analysis (GSEA) also revealed potential phenotypic differences promoted by D463H compared with WT; TGFβ and Myc target signatures are increased, while mutant p53 downregulated genes are suppressed (Fig. 6B); these are pathways broadly implicated in most human tumours, including glioma (*26–29*). D463H also enhanced a signature associated with Cyclin D promotion of cell cycle progression (*30*), which has previously been reported as a downstream growth effector axis for PKCα in glioma (fig. S8A) (*31–33*).–Bioinformatic analysis does not however implicate PKCα D463H in the regulation of common oncogenic signalling pathways, suggesting a non-canonical signalling mechanism underlying deregulation of Myc, p53 and cell-cycle progression (fig. S8A). D463H also enriched signatures for anabolic processes, including protein translation, amino-acid and nucleic acid metabolism, and ribosome biogenesis, all in line with enhanced Myc-driven growth (fig. S8B, C).

**Fig. 6.**
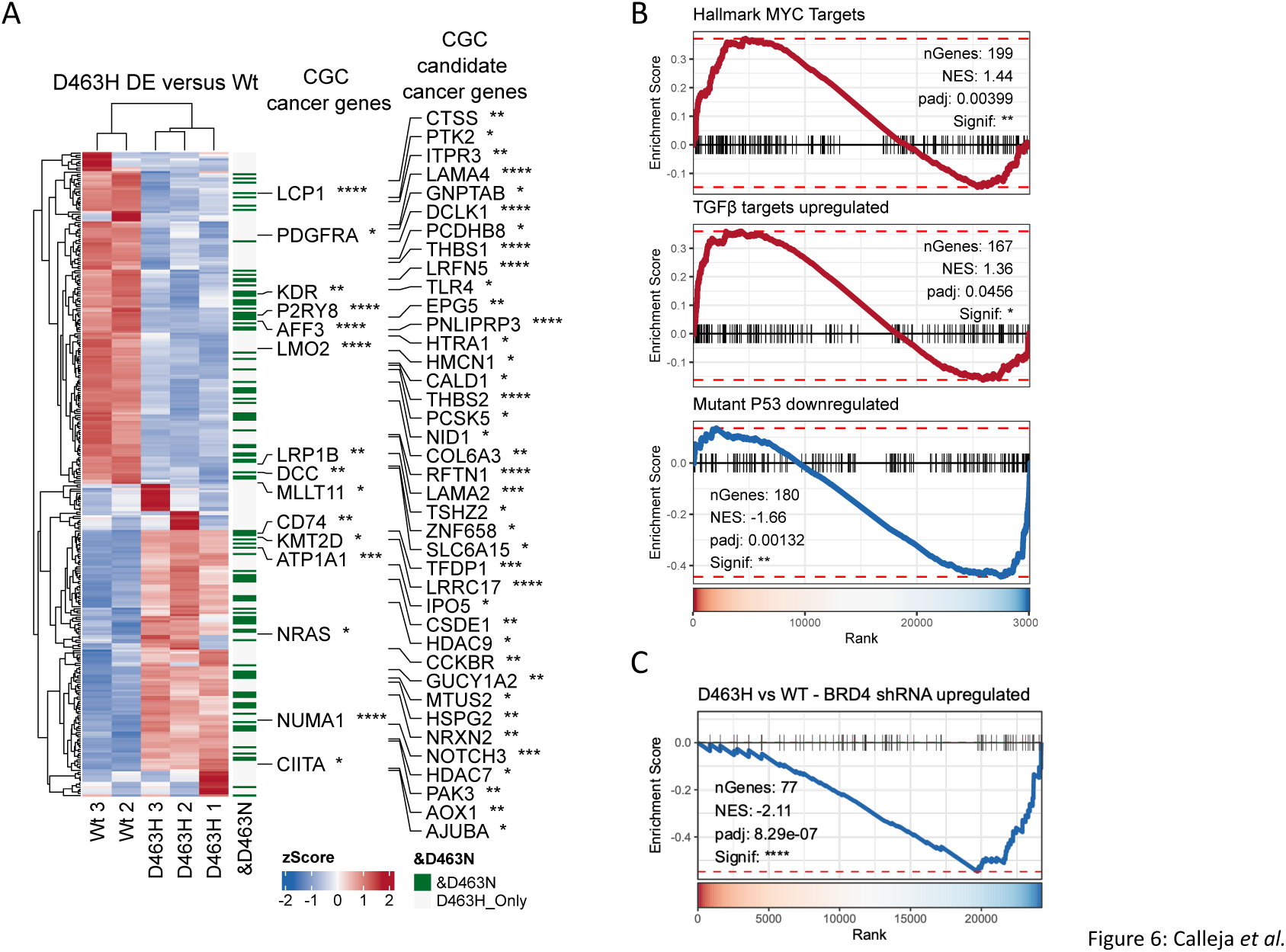
**Differential gene-expression induced by WT-PKC**α **and PKC**α **D463H.** (**A**) Heatmap of genes differentially expressed between PKCα_D463H and WT cells. Right heatmap annotation identifies overlap in significant differentially expressed genes in D463N expressing cells when compared to WT expressing. Known and candidate cancer genes determined by the network of cancer genes (NCG) database of cancer genes are labelled with lines. The significance of the gene differential expression is labelled with the corresponding gene symbol. (**B**) Enrichment plots from GSEA analysis for hallmark Myc pathway and oncogenic TGFβ UP.v1 UP and P53 DN.v1 DN pathways obtained from the molecular signatures database. Ranks were calculated based on the Wald statistic from differential expression results between PKCα D463H and WT cells. Statistical significance was determined using the fGSEA R package. (**C**) Enrichment plot from GSEA analysis for D463H vs WT.

The enhanced interaction of PKCα-D463H with epigenetic regulators provides a potential mechanism driving observed changes to the transcriptome. Intriguingly, the D463H exclusive binder BRD4 has been implicated as an oncogenic regulator of many cancers, including glioblastoma, and widely linked with Myc driven tumourigenesis (*18*). Usefully, Du *et al.* conducted a study examining BRD4 dependent gene-expression in U251 glioblastoma cells (*34*). Using genesets derived from this data and GSEA analysis, we showed that genes upregulated by BRD4 loss are significantly negatively enriched following expression of PKCα-D463H (Fig. 6C). This supports the concept that PKCα-D463H may regulate transcription through targetable epigenetic mechanisms, providing a potential clinical strategy for *PRKCA* mutant choroid glioma, particularly as BET domain inhibitors have been proposed for use in other glioma settings (*35*). We therefore sought to examine the impact of PKC mutant over-expression on sensitivity of U87MG cells to the BET inhibitors JQ1 and AZD5351 (Fig. 7). Over-expression of D463H but not D463N resulted in significantly decreased sensitivity to growth inhibition with both JQ1 and AZD5351 (Fig. 7A, C); D463H increased both the JQ1 and AZD5351 IC50 values by approximately 5-fold relative to parental, WT and D463N expressing cells (Fig. 7B, D). Interestingly, although WT expressing cells consistently showed lower growth inhibition with JQ1 and AZD5351, IC50 values did not significantly change relative to parental cells (Fig. 7B, D).

**Fig. 7.**
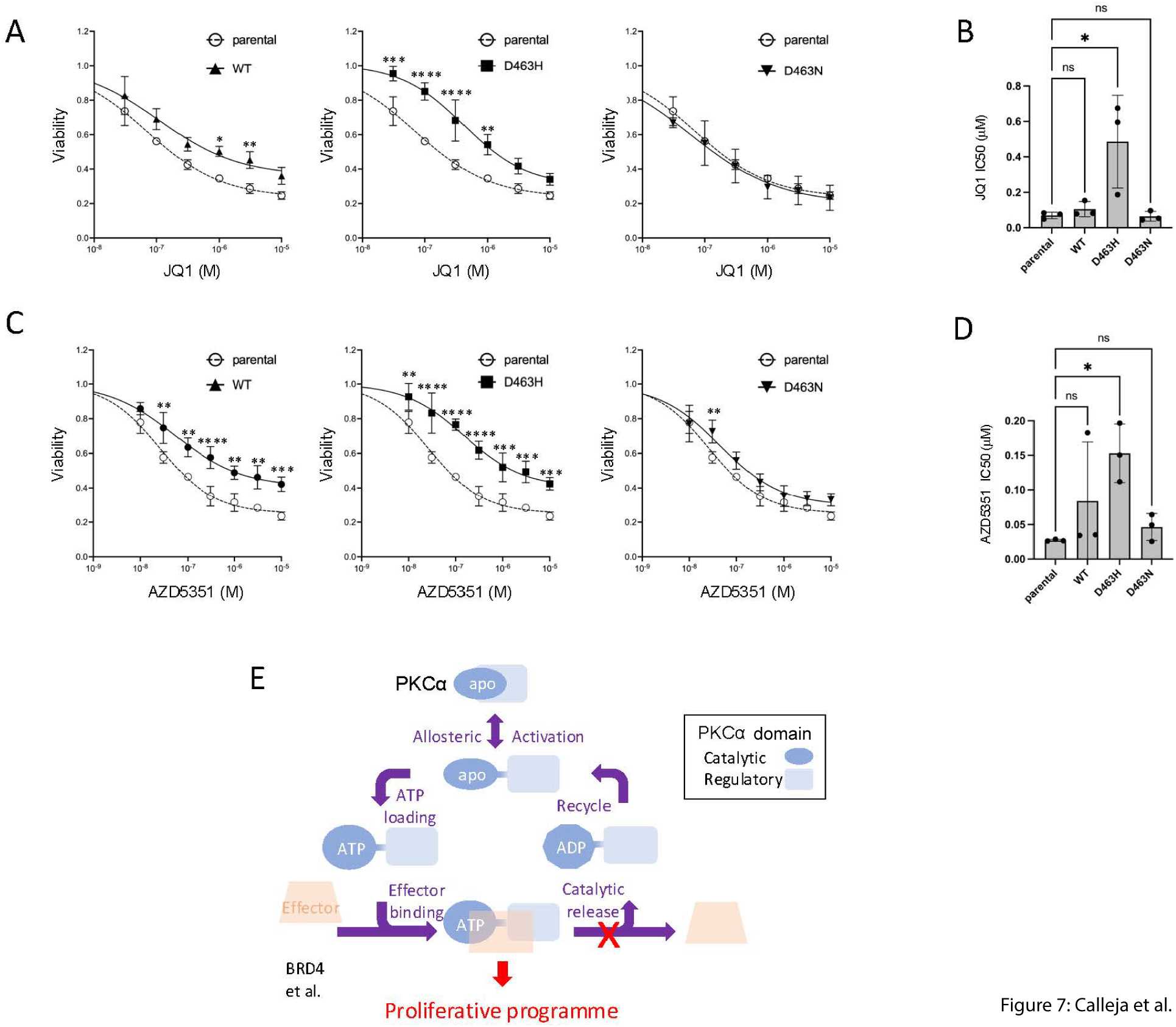
**Resistance to BET inhibitors of PKC**α **D463H mutant expressing cells.** (**A, B**) Inhibition of U87MG WT-PKCα, D463H and D463N expressing stable cell lines inhibited with JQ1. (**C, D**) Inhibition of U87MG WT-PKCα, D463H and D463N expressing stable cell lines inhibited with AZD5351. For clarity of visualisation, each cell lines is separately compared with the parental U87MG cell line. Dose response curves represent the mean viability from three independent biological replicates +/- S.D. Significance was assessed using 2-way ANOVA with Tukey’s multiple comparisons. IC50 values (panels B and D) were calculated separately for each biological replicate and data are presented as the mean +/- standard deviation, with significance assessed by one-way ANOVA compared to the parental control. P<0.05 (*), P<0.01 (**), P<0.001), P<0.0001 (****); n=3. (**E**) Schematic of PKCα and D463H mutant kinase-independent signalling. In the basal catalytically autoinhibited state (Regulatory/Catalytic domains interact), PKCa cannot interact with downstream effector(s); for simplicity this state is indicated as a nucleotide free state (apo) although this is not necessarily the case. On allosteric activation triggering an open conformation, the kinase domain can load ATP, and the ATP-bound state is competent to bind downstream effector(s). In some specific cellular contexts for example in tanycytes and chondrocytes, the formation of a PKCa/effector complex drives a proliferative signal which is switched off through kinase activity-dependent effector release (conformation/stability change). In this model, a tumour promoter such as PMA promotes sustained allosteric activation of the kinase pushing the equilibrium towards a higher effector bound steady state. In the case of the loss-of-activity, gain-of-function D463H mutant that still retains an ATP-bound active conformation, the lack of catalytic activity prevents the normal physiological release of the effector (red cross), sustaining complex formation. The maintenance of the stable PKCa D463H-Effector complex triggers a persistent proliferative output (see text for further discussion).

## DISCUSSION

The early onset chondrosarcomas associated with the heterozygous knock-in of the penetrant chordoid glioma-associated *PRKCA* D463H mutation in mice and the lack of any phenotype associated with the related D463N mutant knock-in (heterozygous or homozygous), provides evidence for an in vivo gain-of-function associated with the D463H mutation. The allied biochemical, interactome and transcriptomic analyses associated with this mutant, suggests that PKCα D463H promotes a non-canonical cell-cycle progression and pro-growth phenotype, which resembles an augmented PKCα driven response. The functional ex vivo evidence indicates that this behaviour is in part driven through epigenetic pathway modulation. Mechanistically, this gain-of-function and loss of catalytic activity is consistent with a working model of effector complex-driven output under negative feedback control of catalytic activity. This is supported by significant modulation of the PKCα interactome for the D463H mutant and structural prediction of a more open kinase conformation (effector competent) (Fig.7E).

The neonatal morbidity of the observed murine chondrosarcomas did not allow us to assess any slow growing chordoid glioma in the brain. The conditional tissue-targeted knock-in of a 463H mutant *PRKCA* allele(s) will be necessary to create a mouse model of this human cancer. Nevertheless, the parsimonious conclusion would be that the driver behaviour of this mutation evident in the early onset chondrosarcomas is conserved in the context of β tanycytes and this has implications for non-surgical intervention. The model suggests that the PKCα D463H mutant preferentially resides in a conformational state, capable of promoting tumour development. This and previous studies indicate that high levels of PKCα, or PKCα retained in an active configuration (for example through sustained effector complex formation) are oncogenic in glioma, but that kinase activity is not required (*12, 36*). This may not be addressable pharmacologically through ATP-competitive inhibitors, indeed this study provides some explanation for why ATP competitive inhibitors of PKCα have shown no benefit in the clinic to date (*1*); to the contrary inhibitors may provoke PKCα output by locking a signalling competent conformer, as promoted by the D463H mutation. We infer that intervention in chordoid glioma would be effective through the removal of mutant PKCα from these tumours. Whilst to date the use of the PKCα-directed antisense agent aprinocarsen has not shown efficacy in other tumour types (see (*37*)), the application of targeted protein degraders offers a powerful alternative route to protein removal (for review (*38*)). For chordoid glioma specifically, this could be developed in a manner selective for the mutant protein, promising a strong therapeutic index.

Targeting oncogenic effectors downstream of PKCα may also provide therapeutic options. The enriched association of a number of epigenetic regulators with PKCα D463H, including BRD4 and MLL1 complex components, certainly warrants closer attention, given that licenced drugs are available for both. Indeed, the enhanced signatures for c-Myc and BRD4, and the functional protection against two distinct BET inhibitors, driven by D463H over-expression in glioma cells, provides strong evidence that this mutant preferentially interacts with the BRD4 axis. As BRD4 and related BET domain family members, and HCFC1, are potent promoters of RNA polymerase II and Myc driven transcription (see for example (*23, 24*)), this could provide a mechanism supporting the gain-of function broad transcriptional upregulation promoted by the D463H mutant compared with WT PKCα. Targeting BET-domain family members has been widely proposed as an approach for suppressing Myc driven tumour growth, including in neuroblastoma and glioma settings (*17, 18, 39–42*). Revumenib which targets MLL1 (also known as KMT2A) has recently been approved for a series of pathway modified tumours (see (*43*)). Preclinical studies to examine the therapeutic potential of such epigenetic targeting drugs in PKCα-D463H mutant choroid glioma are certainly warranted.

Evidence for the underlying molecular mechanisms involved for PKCα have come through multiple avenues contrasting the behaviour of WT and D463H (phenotype inducer) with that of D463N (phenotype neutral) mutants. Comparison of nucleotide binding affinities suggest that D463H preferentially binds ATP over ADP, and with ATP affinity very much higher than the D463N mutant. The affinity of the D463N mutant for ATP would likely predict significant deficit in ATP occupancy at physiological concentrations, which may underly the inability of this mutant to act effectively as a gain-of-function allele. We propose a mechanism where the combination of both ATP binding preference and lack of ATPase activity would benefit an ATP occupied active conformation for D463H-PKCα. Structural modelling suggests both D463H and D463N mutants exist in a PS-out, disinhibited conformation that likely reflects an activated state of the kinase in basal conditions. This ATP-bound effector-competent conformation of the D463H mutant in the absence of activators is further supported by the Turbo-ID profile. This marks D463H out as a privileged gain-of-function mutation analogous to gain-of-function small GTPase mutations which retain unhydrolyzed GTP (Fig. 7E) favouring the effector binding conformers.

Chondrosarcomas are a group of bone malignancies thought to derive from chondrocyte and chondroblast progenitor cells (see (*44*)), and are characterised by the production of cartilaginous extracellular matrix, with a low percentage of dividing cells, as reported here for D463H mice hindlimbs (Fig. 1C). The Archs4 database (https://maayanlab.cloud/archs4/) indicates that PKCα is highly expressed in chondrocytes and is predicted to be linked to an abnormal chondrocyte physiology phenotype in mice (MP0009780). There is also involvement of PKCα in the regulation of extracellular matrix organization (GO:1903055) chondrocyte proliferation (GO:0035988) and cell adhesion mediated by integrins (GO:0033627) (https://maayanlab.cloud/archs4/gene/PRKCA). In concordance, our GSEA analysis of the canonical pathways also showed a decrease in the extracellular matrix organisation, osteogenic collagen synthesis and integrin signalling associated with PKCα D463H expression. This suggests some canonical effects of this specific PKCα D463H mutation may be shared despite the distinctive cellular contexts (Fig. S8B, C). In accordance with the early onset of the phenotype in our heterozygous mice, cartilage formation is one of the earliest morphogenetic events during embryonic development (for review see (*45*)).

Chondrocytes translate mechanical forces in various cellular responses to regulate cartilage homeostasis and function (*46*) and the pathogenesis of bone and cartilage are linked to alterations in these mechanotransduction pathways (for review see (*47*)). It is important to note that mechanical strains occur specifically in the hindlimbs (as opposed to the forelimbs) that support the whole mouse weight, which might explain the localised phenotype seen in our D463H mutant mice, with no evidence of forelimb-associated pathology. Notably, PKCα was shown to be important in the mechanotransduction of the signal downstream of integrins in chondrocytes (*46*) through a role for the PKCα/ERK-1 axis (*48*). Goode *et al.* had shown that ERK signalling was upregulated in chordoid glioma, likely associated to the presence of D463H mutation in these tumours (*5*) and here functional GSEA pathway enrichment in the glioma cell line showed also an enrichment of the PDGF_ERK up regulated geneset on expression of the PKCα D463H mutant (Fig. S8A). Altogether the results suggest that the formation of chondrosarcomas in the hindlimbs of D463H mutant mice might be due to PKCα D463H driven dysregulation of the mechanotransduction signals involved in cartilage homeostasis, through pathways we anticipate would be conserved in the context of chordoid gliomas.

It is notable that various kinase inactive PKCα mutants have been employed in different studies to examine their dominant negative impact on physiology, our results suggest that the preconceived expectation of their impact needs reassessing. The likelihood that in some contexts, a protein-driven pathway may indeed be negatively influenced by catalytic activity should now be considered. For example, it has been shown that expression of the inactive PKCαK376R mutant in haematopoietic progenitor cells which are then differentiated down a B-Cell lineage, triggers the formation of a B-CLL phenotype (*49*). The original interpretation promoted a suppressive function in B-cells, re-evaluation in light of the results here might lead to a rather different conclusion. Generally, the penetrance of catalytic activity independent properties of PKCα and indeed other members of this family and the wider kinome would merit broad consideration. In the context of pseudokinases we readily accept the importance of allostery and the associated influence of ATP binding on kinase domain conformation (see (*50*)). The implication of the current findings is that such properties can be ascribed to active kinases and moreover that activity may act negatively as well as positively on kinase effector output(s).

The evidence presented provides a paradigm shift in our appreciation of PKCα action and questions our understanding of the wider kinome. Unravelling the conformation-dependent but kinase-independent proximal targets and associated downstream programs elicited by this PKCα mutant, will provide insights that will no doubt guide the development of innovative interventions.

## MATERIALS AND METHODS

### Antibodies and Reagents

Phospho-PKC Substrate Motif [(R/K)XpSX(R/K)] MultiMab™ Rabbit mAb mix (#6967), Phospho-(Ser) PKC Substrate Antibody (#2261), Phospho-PKC (pan) (gamma Thr514) Ab (#9379), Phospho-PKCα/β II (Thr638/641) Ab (#9375), Phospho-PKC (pan) (βII Ser660) Ab (#9371) and PKCα Ab (#2056) were from Cell Signaling Technology. IRDye800CW Streptavidin (926–32230), anti-rabbit IRDye800CW (925–32211), anti-mouse IRDye680RD (926–68070) and Odyssey Blocking Buffer (TBS) were from Li-COR Biosciences. Anti-HA tag Ab [HA.C5] (AB18181) and Anti-Myc tag [9E10] (ab32) were from Abcam. SYPRO orange protein gel stain (S5692) was from Sigma-Aldrich. Mouse monoclonal [9F9.F9] to GFP (Ab1218) Abcam. InSolution phorbol 12-myristate 13-acetate (PMA) was from Merck. FuGENE^®^HD (E2312) was from Promega. cOmplete™, EDTA-free Protease Inhibitor Cocktail (12326400) was from Roche. Streptavidin-HRP conjugate (GERPN1231) from Sigma-Aldrich. Lipofectamine RNAiMAX Transfection Reagent (13778150) Life Technologies LTD. HA-probe (F-7), monoclonal antibody (sc-7392). Complete mini EDTA free (4693159001) from Sigma-Aldrich. RNeasy Mini Kit (74104) Qiagen LTD. QuantiTect SYBR Green RT-PCR Kit (204243) Qiagen LTD. ADP-Glo™ kinase assay (V6930) Promega. In-Fusion cloning kit was from Takara (638948). Phosphatase inhibitors set II (524625) and IV (524628) were from Millipore.

### CRISPR/Cas9 genome editing for *Prkca* D463H knockin mutation in mice

All animal studies were compliant with UK Home Office regulations and carried out under license. Prkca protein kinase C, alpha [*Mus musculus* (house mouse)] -Gene ID: 18750. The CRISPR-Cas9 reagents were delivered directly into the mouse zygote to derive a mutant mouse carrying targeted genetic modifications. The single guide sgRNAs which provide target specificity for D463>H mutation in exon 13 on chromosome 11 were Guide RNA 1 and Guide RNA 2. The protocol uses ZEN, 1.2uM/6uM/300ng/μL, N, 2c, Repair – x1 ssODN, 75, Guide1 and ZEN, 1.2uM/6uM/300ng/μL, N, 1c, Repair – x1 ssODN, 75, Guide2 respectively was used for 107 embryo injections in total. The Cas9 nuclease specificity that creates the DNA double-strand break is dictated by the PAM motif a trinucleotide sequence of NGG, directly adjacent to and continuous from the 3’end of the protospacer sequence of the noncomplementary strand. The donor oligonucleotide or plasmid carrying the intended mutation flanked by sequences homologous to the target site were injected with the guide RNAs (table S10).

### Mouse cohorts

Mouse strains with D463H mutation are: IRCP 1, 3 and 4, mosaic mice C57BL/6Jax (table S1); PPBR 5-10, Het mice C57BL/6Jax x C57BL/6Jax cross (table S2) ; PPBY 1, 4 and 5: Het mice C57BL/6Jax x CD-1 cross (table S3); PPBY 6 and 7, Het C57BL/6J x CD-1 cross in trio with CD-1 (table S4). The second round of CRISPR for the D463H mutation are: IRCP 8, mosaic mice with C57BL/6Jax background; IRCP 20 mosaic mice with F1(B6 x CBA) background; PPCE 1 and 2: Het mice C57BL/6Jax x CD-1 cross (table S5); PPCE 7 and 8: Het mice F1(B6 x CBA) x CD-1 cross (table S5); Mouse strains with D463N mutations: IRCY 3 and 6, mosaic mice with C57BL/6Jax background (table S6); IRCY 11, 12 and 13, Het mice C57BL/6Jax x C57BL/6Jax cross (table S7); PPBZ 2 and 3, PPCA 1, PPCA 11 and PPCB 1 are Het mice C57BL/6Jax x C57BL/6Jax cross (table S8); PPCA 4 are Het mice C57BL/6Jax x CD-1 cross; PPBZ 7 are Het mice C57BL/6Jax x F1(B6 x CBA) cross (table S8). PPBZ 8 and PPCB 12 are homozygous mice C57BL/6Jax x C57BL/6Jax cross (table S9). No phenotypic variation was observed between the sexes.

### Histopathological Examination

Four µm-thick, formalin-fixed, paraffin-embedded (FFPE) sections were stained with haematoxylin & eosin (HE) and examined by two board-certified Veterinary Pathologists (ASB & SLP). Histopathological assessment was performed blind to experimental grouping using a light microscope (Olympus BX43). Tissue sections were examined individually and in case of discordance in diagnosis a consensus was reached using a double-headed microscope. Sections were histopathologically assessed using the INHAND guide for non-proliferative and proliferative lesions of the rat and mouse skeletal tissues (*51*).

### Recombinant protein production in insect cells

For the generation of the baculovirus, Sf21 insect cells were co-transfected with the pTRI-EX vector containing PKCα and FlashBAC™ backbone baculovirus vector. The viruses were amplified by a series of 0.22 µm filtered viral supernatant infection of Sf21 cells. The extraction of the recombinant proteins was done by resuspending the spun cells in wash buffer: 50 mM Tris-HCL pH 7.5, 150 mM NaCl, 1 mM EDTA and 1 mM DTT containing 1% Triton X-100 with addition of protease inhibitors for lysis. The centrifugated lysate supernatant was put on Glutathione-Sepharose beads 4B and left to bind for 2h on roller at 4°C and the beads washed 3 times with the wash buffer. For elution of the GST-proteins, the beads were resuspended in wash buffer containing 20 mM Glutathione reduced (pH 7.5) and left over/night at 4°C in slow motion roller/rotisserie. For 3C or TEV cleavage, the beads were re-suspended in wash buffer with 3C-GST or TEV-GST proteases and the cleaved recombinant proteins were recovered by centrifugation. For larger scale protein extraction, the same protocol was used, and the proteins were spun at 4,000 rpm at 4°C down to 500µl using a Vivaspin 6 concentrator (30,000 cut off for full-length PKC and 10,000 cut off for kinase domain). The proteins were purified on Superdex S200 AKTA. The correct expression of GST-PKCα full length and kinase domain was confirmed by SDS-PAGE Coomassie gels and their phosphorylation by Western blot using anti-phosphospecific antibodies.

### Pull down GFP-DARPin and kinase assay

All cell lines were authenticated by short tandem repeat (STR) profiling and Mycoplasma screened by a PCR-based approach by Cell Services at The Francis Crick Institute. U87MG transiently transfected with GFP-PKCα and mutants were solubilised in lysis buffer (20 mM Tris, pH8; 130 mM NaCl supplemented with 1% Triton X-100). GFP-PKCα constructs were pulled down using DARPin (Designed Ankyrin Repeat Proteins) agarose beads directed against GFP and subjected to a kinase assay using 20 mM TRIS pH8, 2 mM DTT, 100 nM Calyculin A, 10 mM MgCl2, 1 mM ATP supplemented with PKC activators: 400 µg/ml PtdSer, 1 µM PMA, 300 µM CaCl2. GFP-PKCα and associated substrates phosphorylation were detected by western blot using anti PKCα phospho-substrates and anti GFP antibodies.

### Fractionation assays

After being untreated or treatment with 500nM PMA for 4h or 24h, U87MG cells were solubilised in 20 mM Tris, pH8, 130 mM NaCl, 1% Triton X-100 supplemented with protease inhibitors for 15 min at 4°C. the cells were scraped and put in an Eppendorf and centrifuged at 16,000g for 10 min at 4°C. LDS/DTT (4X) was added to the supernatant (soluble fraction) and the pellet (Insoluble fraction) was diluted in (2X) LDS/DTT for loading on an SDS-PAGE gel. The proteins were transferred onto a PVDF membrane and the amount of PKCα in each condition was detected by western blot using an anti-HA antibody. The phosphorylation of PKCα hydrophobic motif over total protein was detected using an antibody anti phosphor-Serine 657 concomitantly with an anti-HA antibody (Li-COR).

### RNA sequencing and analysis

Stable U87MG cell lines expressing Myc-PKCα WT, D463H and D463N and parental control cells were grown and plated on a 6-well plate to isolate mRNA. The RNA extraction was done as per manufacturer protocol (RNeasy kit from Qiagen) on about 1 to 1.5 M cells. The mRNA from 3 independent experiments were sequenced and analysed for each construct by the high throughput sequencing facility technology platform at the Francis Crick Institute. Differential expression analysis was conducted using the R package DESeq2 (*52*) and batch correction of counts using the R package limma (*53*). Bioinformatics was performed using bespoke R scripts. Known and candidate cancer genes were obtained from the Cancer Gene Consensus (CGC, COSMIC) (*54*). Overrepresentation analysis was performed using the R package clusterProfiler (*55*) using ontologies obtained from the Gene Ontology (GO) resource (*56*). Geneset enrichment analysis (GSEA) was performed using the R package fgsea (*57*) using genesets curated from the molecular signatures database (MSigDBv7.5.1). The BRD4 shRNA upregulated geneset was curated by extracting the top 100 upregulated genes with BRD4 shRNA treatment in U251 cells previously published (GSE97791, GEO). GitHub link: Custom R scripts used to analyse and plot data are publicly available at https://github.com/jackchenry/PKCa_Calleja-et-al.

### TurboID experiments

Stable U87MG cells expressing constitutively 3xHA-TurboID-PKCα fusion constructs WT and mutants were plated on a 6-well plate at 500,000 cells/well. Biotin (500 µM) was added the day after for 10 to 30 min. The cells were then washed extensively to remove the free biotin, lysed with RIPA buffer (50mM Tris-HCl, pH 7.4, 150mM NaCl, 0.5% deoxycholic acid, 1% NP-40, 1mM EDTA) supplemented with 0.1% SDS and protease inhibitors cocktail (cOmplete™ mini from Roche) and pulled down with neutravidin agarose beads (Thermo Fisher Scientific). After 3x washes using the lysis buffer at 4°C the beads were recovered in sample buffer and the proteins separated by SDS-PAGE. After transfer onto PVDF membrane the biotinylated proteins were detected by western blot using streptavidin labelled with an IRDye 800 (LI-COR/Odyssey detection system). For simple detection of protein expression upon various treatments the stable cell lines were directly lysed by addition of Nu-PAGE LDS sample buffer (Thermo Fisher Scientific) supplemented with DTT (1/10e) separated by SDS-PAGE and detected by western blot with streptavidin-IRDye800. In parallel the expression of the PKC constructs was verified by western blot with an anti-HA antibody and a secondary IRdye680 (red pseudocolour).

### Thermal shift assay

The thermal shift assay is an indirect measurement of protein stability. Upon increase in temperature proteins unfold allowing the binding of the fluorescent dye SYPRO orange. The increase in fluorescence versus temperature is recorded in real time (melting curves). The stability of PKCα and mutants was measured in the apo form or when stabilized with addition of increasing amount of ATP or ADP (15 dilutions ranging from 0.1µM to 5 mM). On a 384- well plate each condition was acquired in triplicate, 0.4 to 0.7 µg of recombinant protein PKCα WT or mutants was added without or with a serial dilution of ATP or ADP ranging from 0.01 mM to 5mM. The proteins and nucleotides were diluted in the thermal shift buffer containing 50 mM HEPES, 150 mM NaCl, 1 mM DTT supplemented 10mM MgCl2. SYPRO orange (1:1000 in thermal shift buffer 1X) was then added to the mix. The plate was placed in a QuantStudio7 flex RT-PCR with Melt curve standard program gradually increasing the temperature from 15°C to 75°C with an incrementation of 0.5°C/s. The melting curves (ascending part of the curve) were fitted with a non-linear sigmoidal or a biphasic fit using GraphPad/Prism software at each concentration of ATP or ADP to determine the melting temperatures (Tm). The stabilization effect (βTm) of the ligand for each protein was subsequently obtained by subtracting Tm[ligand] – Tm[apo] at each concentration of ATP or ADP. The EC50 were obtained by plotting the βTm versus [ATP] or [ADP] and fitting the curve with GraphPad/Prism.

### PKC recombinant protein labelling via amine coupling with NHS RED dye for MST analysis

The labelling of the PKCα full length recombinant proteins was done using Monolith protein labelling kit RED-NHS 2^nd^ generation (Nanotemper) according to manufacturer’s protocol. Briefly, the recombinant proteins were first buffer exchanged into 50mM Hepes, 150 mM NaCl buffer prior to labelling to exchange from the original buffer containing Tris. Proteins and dye were incubated for 30 mins at room temperature. The excess dye was removed using a spin column. The proteins were eluted in 450 µl of MST buffer 50mM Hepes, 150 mM NaCl, 10mM MgCl2 and fresh 1µM DTT. The aim was to obtain a ratio of dye to protein in the order of 0.3 to 1 for the MST experiments.

### Microscale thermophoresis (MST)

PKCα full length recombinant proteins labelled with NHS RED and diluted in the MST reaction buffer were mixed with ATP at various concentrations. For each assay, 12 x 20 µl samples containing the recombinant proteins at a final concentration of 50nM or 100nM as used. 20 µl of ATP was added to the reaction in a 2-fold serial dilution up to final max concentration of 30 µM or 60µM for mutant D463N. Samples were transferred to premium Monolith NT.115 capillaries. Experiments were run with a LED power of 20 % and MST power of 20 %, at 25 °C, with a 5s-20s-5 run. Run on Monolith NT.115 (Nanotemper technologies) machine, using NT Control v.2.2.1 (Nanotemper Technologies) software and analysis performed in MO Affinity Analysis v.2.3 (Nanotemper Technologies) software. The Kds were determined by fitting the data to a non-linear regression using a quadratic binding model.

### Cloning of the PKCα constructs

pEGFP-C1-PKCα (GFP-tagged human PKCα) is an in-house construct (The Parker laboratory at the Francis Crick Institute). Constructs were created by InFusion® cloning in vectors cut with EcoRI. pCDNA3-Myc-PKCα was performed by cloning PCR amplified PKCα with an N-terminal Myc tag using the oligos Myc-PKCα_sense and _anti. pBABE puro-Myc-PKCα was done by cloning of PCR amplified Myc-PKCα using the oligos Myc-PKCα_for and _rev. pBABE puro-3xHA-TurboID-PKCα was done by cloning PCR amplified 3xHA-TurboID from the vector Addgene 3xHA-TurboID (#107171) upstream of the 6xGly-PKCα PCR product using the oligos 3xHA-TurboID_for and _rev and 6xGly-PKCα _for and _rev. pTriEX6-GST-PKCα insect cells expression construct was done by cloning the PCR product of full length PKCα into the vector cut with EcoRI-BamHI, using the primers GST-3C PKCα_for and _rev. pTriEX6-GST-PKCα kinase domain was done by introducing a TEV cleavage site (E-N-L-Y-F-Q-|-G/S, cleavage between Q and G/S) in the hinge region just after the residue 321 of full length PKCα in the pTriEX6-GST-PKCα construct. PKCα regulatory domain, a fragment containing the TEV site, and PKCα kinase domain with the C-terminal were PCR amplified separately. The three PCR products were introduced into pTriEX6-GST-3C cut with EcoRI-BamHI. The oligos used for the 3 PCR amplifications are listed in table S11.

### Stable Cell lines

To generate stable polyclonal cell lines expressing Myc-PKCα for the RNAseq experiments, U87MG cells were plated in a 6-well plate at 300,000 cells per well and transfected 24h later with pBABE puro-Myc-PKCα WT or mutants. The cells were then selected with puromycin until the control non-transfected cells were all dead (approx. one week). To prepare stable polyclonal cell lines expressing 3xHA-TurboID-PKCα WT and mutants for the identification of protein partner patterns, cells were transiently transfected with pBABE puro-3xHA-TurboID-PKCα and mutants then selected with puromycin until the control non-transfected cells were all dead (approx. one week).

### PMA-induced PKCα degradation experiments

U87MG cells expressing transiently GFP-PKCα or Myc-PKCα WT and mutants were treated with 500nM PMA for various durations up to 24h. The time course decrease of PKCα priming site phosphorylations were detected by western blot using anti phospho-specific HM, turn motif and activation site antibodies, and the expression of each protein determined with anti GFP or anti Myc antibody.

### Kinase activity of recombinant PKCα full length or kinase domain

After determination of the optimal PKCα or PKCα kinase domain concentration (linear range of protein v activity), the Km for ATP was determined by dose response of ATP. 50 ng of recombinant enzyme was mixed with 25µM ATP (approx. Km) and with activators in kinase buffer in the presence or absence of substrate either 0.2 mg/mL peptide substrate (PPSS) or 0.5 mg/mL of Protamine sulphate (no activators required). The kinase buffer (1X) was prepared as follow: 40 mM Tris/HCL pH 7.5, 20 mM MgCl2, 0.1 mg/mL BSA and 50 µM DTT. The activators 400 µg/mL of phosphatidylserine (PtdSer) in 1% TX-100, 1 µM PMA and 300 µM CaCl2 were added only with the PPSS peptide (Ac-RFARKG**<u>S</u>**LRQKNVH-CONH2). The reactions were performed as specified by the manufacturer’s protocol (ADP-Glo™ -Promega).

### Preparation of the samples for immunoprecipitation and mass spectrometry

U87MG cells were plated at 500, 000 cells / well on a 6-well plate. The following day the cells were transfected with 2µg / well of Myc-PKCα WT or D463H or D463N mutants. On day 4, the cells were lysed in 250 µl / well of lysis buffer containing 20mM Tris pH 8.0, 130mM NaCl, 1% Triton X-100, 1mM DTT, 10mM NaF with protease and phosphatase inhibitors (ROCHE). 3 wells were used per condition. The immunoprecipitation was performed using Myc-trap®Agarose (ChromoTek). After 3 washes with lysis buffer without inhibitors, a kinase assay was done on the IPs (with Mg-ATP) in a vol. of 50µl per condition. The supernatant was recovered 50µl and 20µl of LDS 4X/DTT was added. The beads were recovered in 50µl of LDS 2X. The proteins were separated after a short run by SDS-PAGE on a 10 % Bis-Tris gel. The gel was stained with Coomassie blue, and the bands were cut and processed for mass spectrometry. A Western blot (Li-COR Odyssey®) with total Myc tag and phospho-PKC substrates ab was performed in parallel to control the success of the kinase reaction and of the Myc-PKCα pull down.

### Label free quantitation mass spectrometry and data analysis

Proteins were digested with trypsin, and peptides extracted according to the protocols established by the Proteomics STP lab at the Francis Crick Institute. Peptides were separated on a 50 cm, 75µm I.D. EasySpray column over a 120 min gradient and eluted directly into an Orbitrap Fusion Lumos mass spectrometer. Xcalibur software was used to control the data acquisition. The instrument ran in Data Dependent Top Speed mode with cycle time of 3s. The most abundant peptides with charge states 2-4 were selected for MS/MS. Data processing was performed by the MaxQuant bioinformatics suite using the LFQ algorithm and “match between runs” option. The Homo sapiens protein database was searched. Any peptide identified from the control samples was removed from analysis. Data was plotted using custom R scripts. Known PKCα interaction data was curated from the STRING and BioGRID databases. Further data analysis and presentation were performed with the Perseus software.

### STRING analysis

STRING analysis (*19*) was conducted on proteins recovered in PKCα immunoprecipitates. Any proteins recovered in control precipitates were excluded, as were those only recovered in a single sample. For analysis of proteins preferentially associating with the D463H mutant, proteins showing greater than a 0.5 log FC enrichment in the D463H samples over the WT samples were considered hits. This list also includes all D463H exclusive binders. STRING analysis was set at varying levels of confidence with only direct binders included in the query list (no shell); text mining, experiments, databases, gene fusion and co-occurrence are included as potential interaction sources.

### Cell viability assays

U87MG parental, WT-PKCα, D463H and D463N expressing stable cell lines were plated in 96 well plates in 100ul of complete DMEM with a minimum of five technical replicates. 24h later cells were treated with either JQ1 or AZD5351 in a final concentration of 0.1% DMSO (Vehicle). A no growth control well was also treated with sodium azide (0.1%). Cells were grown for a further 72 hours prior to the addition of 0.5 mg/ml MTT for 3 hours. Formazan crystals were solubilised in DMSO, and absorbance was measured at 570nm using a colorimetric plate reader (BMG Spectrostar). Cell viability was calculated relative to Vehicle and azide controls. All experiments were conducted as a minimum of three biological replicates. For curve fitting and IC50 calculations, [inhibitor] vs response analysis (Non-linear fit, variable slope (four parameters) was conducted using Prism software.

### In silico mutant predictions

AlphaFold 3 was used to predict the structure of full-length (fl) PKCα WT, D463H or D463N, in presence of 1x ATP, 2x Mg^2+^ and 4x Zn^2+^. Twenty seeds (1–20) were used, generating a total of 100 models per protein construct. To classify predictions into PS-in or PS-out conformers, the PAE matrix was used. The mean score of a region 1 with X between [0, 20], Y between [350, 650] (PS-kinase domain interaction) was compared to a region 2 with X between [200, 250], Y between [350, 650] (C2-kinase domain interaction). If the mean error score of region 1<2, the observed conformation was considered PS-in, whereas if the mean score of region 1>2, the observed conformation was considered PS-out. To predict the biophysical effect of D364 substitutions ddMut (*58*) was used with the top-scoring fl-PKC WT model as input.

### Statistical Methods

Statistical analysis of data was performed using t-tests where comparisons are made for groups where there is equal variance. Where we are comparing values relative to controls, we have employed a one way anova. A minimum of n = 3 independent experiments was used (n is indicated in figure legends). Dose response curves represent the mean viability from three independent biological replicates +/- S.D. Significance was assessed using 2-way ANOVA with Tukey’s multiple comparisons. IC50 values were calculated separately for each biological replicate and data are presented as the mean +/- standard deviation, with significance assessed by one-way ANOVA compared to the parental control. P<0.05 (*), P<0.01 (**), P<0.001), P<0.0001 (****).

## Supplementary Materials

### Supplementary Figures and Tables

fig. S1. Forelimbs and cranial analysis in heterozygous PRKCA-D463H knockin mice.

fig. S2. Recombinant PKCα constructs expression and phosphorylation.

fig. S3. Thermal shift assay and microscale thermophoresis.

fig. S4. Similar regulation of WT-PKCα and D463H mutant constructs upon PMA treatment.

fig. S5. TurboID-PKCα biotinylation of PKCα and mutants.

fig. S6. Structure predictions for mutant forms of PKCα.

fig. S7. PKCα D463H is a gain-of-function mutant.

fig. S8. GSEA analysis of differential oncogenic and canonical pathways between WT-PKCα and D463H.

table S1. *Prkca*-D463H mosaic mice obtained from the first round of CRISPR/Cas9 genome editing.

table S2. *Prkca* D463H Het mice obtained from IRCP4.1c mosaic mice by timed mating.

table S3. *Prkca* D463H Het mice obtained from IRCP4.1c mosaic mice by IVF of CD-1.

table S4. *Prkca* D463H cross for Het mice with PPBY4.1c in trio with CD-1.

table S5. *Prkca* D463H Het mice obtained from 2 mosaic mice from second round of CRISPR/Cas9 genome editing.

table S6. *Prkca*-D463N control mosaic mice obtained from CRISPR/Cas9 genome editing.

table S7. *Prkca* D463N control Het mice obtained from three mosaic mice from CRISPR/Cas9 genome editing.

table S8. *Prkca* D463N control second round of cross for Het mice.

table S9. *Prkca* D463N control cross for Homozygous mice.

table S10. Guide RNAs for knock-in studies.

table S11. Oligonucleotide sequences for construct cloning.

table S12 Interactors PKCα WT, PKCα-D463H or PKCα-D463N (Excel file). table S13. Exclusive PKCα mutant D463H interactors.

## Acknowledgments

We would like to thank Dr Aengus Stewart former head of the bioinformatic laboratory at the Francis Crick Institute for his support in the supervision of the RNA sequencing data analysis and Dr Ian Rosewell former head of the Gene modification laboratory at the Francis Crick Institute for his invaluable input and advice on the strategy, design, and analysis of the genetically modified (CRISPR) mouse cohorts. We also would like to thank the BRF team and specifically Samuel Cooper and Nicolas Chisholm for their commitment and wonderful work in the challenging maintenance and care of the knock-in mice cohorts. A big thank you to the Science technology platform deputy head Dr Svend Kjaer and associates Dr Annabel Borg and Dr Roger George for critical help in the design and production of the recombinant proteins.

## Funding

The study was funded by the Francis Crick Institute, which receives its core funding [CC2068 and CC2140] from Cancer Research U.K., U.K. Medical Research Council and the Wellcome Trust. AJMC is supported by Cancer Research UK Centre Grants to Barts Cancer Institute (C355/A25137) and the City of London Centre (C7893/A26233). AJMC, JCH and JS are supported by MRC grant MR/X018997/1.

## Author contributions

Conceptualization: PJP, AJMC

Investigation and analysis: VC, JCH, MC, KR, SB, NA, SV-B, TS, AS-B, SLP, JS

Funding acquisition: AJMC, NQM, PJP

Writing – original draft: VC, PJP, AJMC

Writing – review & editing: all authors

## Competing interests

There are no conflicts of interest for any authors.

### Data and materials availability

Proteomics data is available at data deposited at PRIDE accession number PXD049959 and RNAseq data is available at the GEO repository accession number GSE269531. All other data required to evaluate the conclusions of the paper are presented in the text, figures, legends, and supplementary materials. Materials are available through the corresponding authors.

## Notes

https://www.ncbi.nlm.nih.gov/geo/query/acc.cgi?acc=GSE269531

### Competing Interest Statement

The authors have declared no competing interest.

