## Supplementary Material for "The penetrant chordoid glioma PRKCA mutation is an oncogenic gain-of-function kinase inactivation eliciting early onset chondrosarcoma in mice"

### Supplementary Materials

### Supplementary Figures

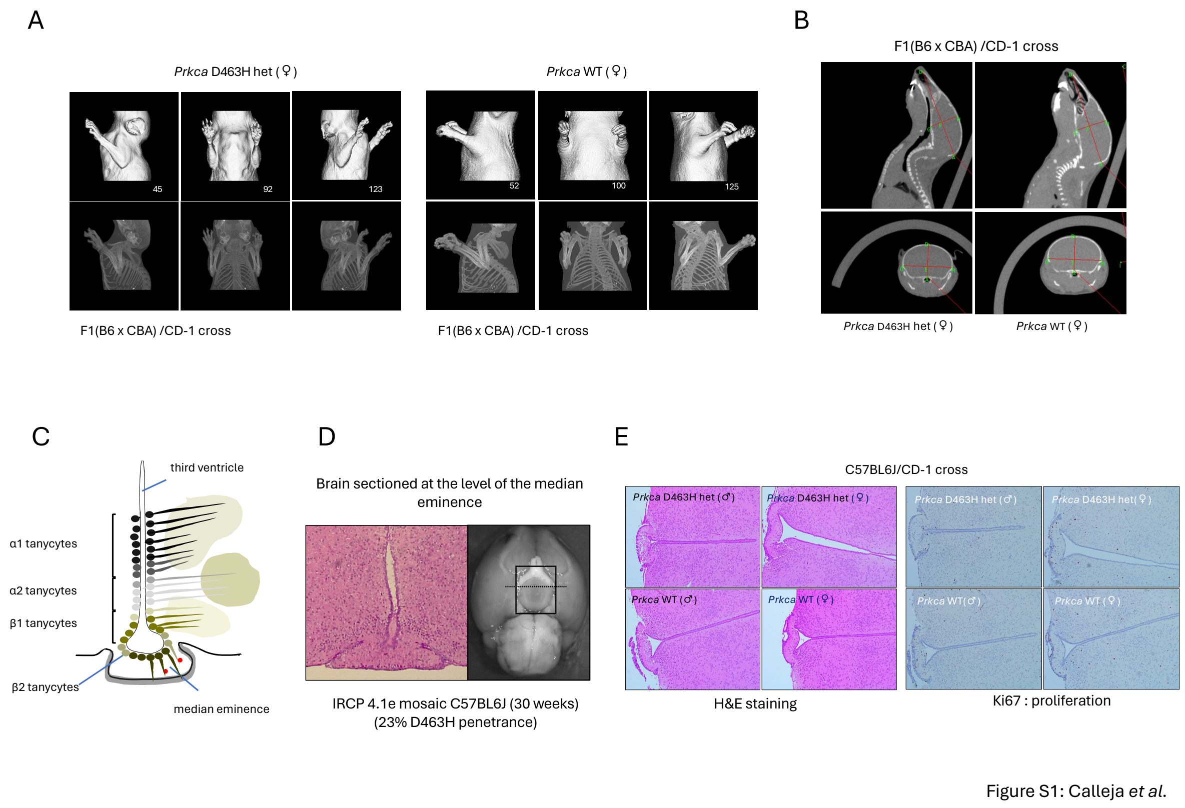


**Fig. S1. Forelimbs and cranial analysis in heterozygous PRKCA-D463H knockin mice.** (**A**) CT-scan of the forelimbs of a D463H mouse compared to WT. There is no visible growth of cartilage in the forelimb's due to the heterozygous mutation. (**B**) CT-scan of a representative head of a D463H Het mouse compared to WT. There is no visible change in the morphology when related to body weight. (**C**) Schematics of the brain's median eminence and the different type of cells. (**D**) no morphological change can be seen by H&E on a 23% D463H mosaic brain's median eminence of a 30 week old mouse. (**E**) H&E and Ki65 median eminence staining of D463H het (male and female 2 weeks old) mice. No morphological changes or proliferation can be detected in comparison to the WT littermates.

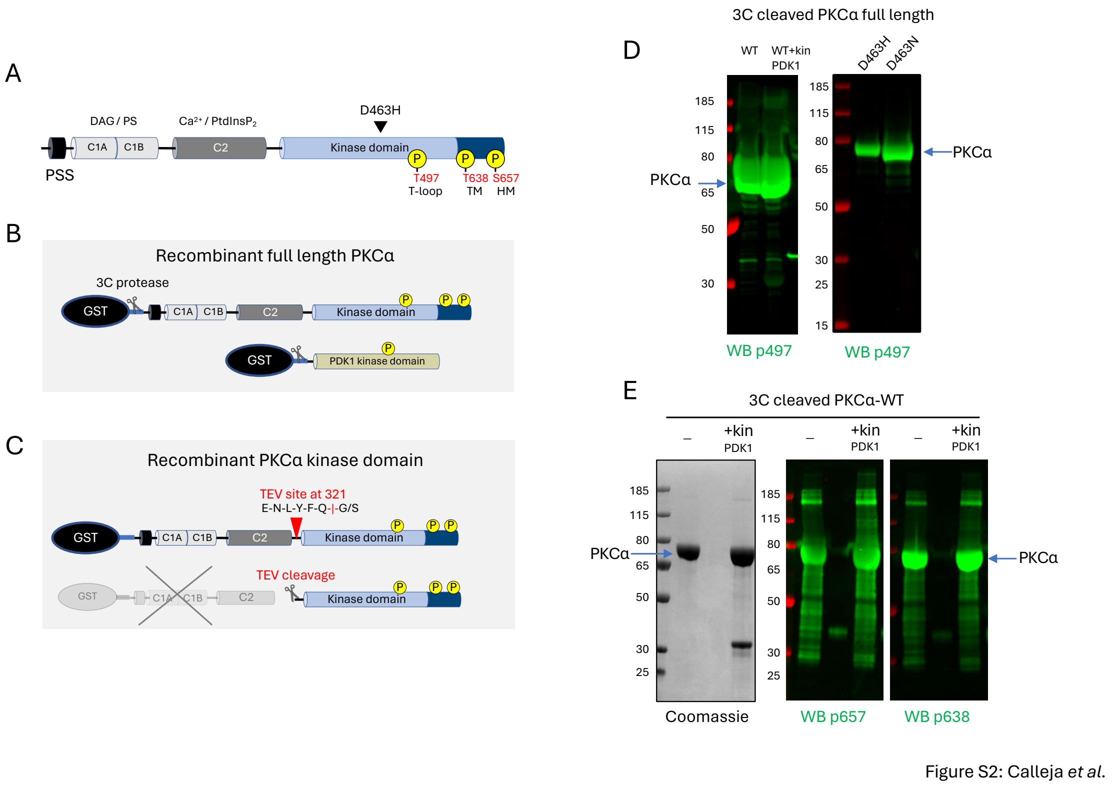


**Fig. S2. Recombinant PKCα constructs expression and phosphorylation.** (**A**) representation of the PKCα domains sequence and phosphorylated activation sites in the kinase and C-terminal domains (T-loop or activation loop; TM: turn motif; HM: hydrophobic motif). C1A and C1B binding sites for diacylglycerol (DAG or phorbol ester like PMA) and anionic phospholipids (1,2-*sn*-phosphatidyl-L-serine) at the plasma membrane, regulate PKCα kinase domain activation. The activation of the kinase is also dependent of the C2 domain through calcium-dependent binding to inositol bisphosphate ($PtdInsP_2$) at the plasma membrane. The pseudo substrate site (PSS) is shown in N-terminus of the sequence. The D463H mutation is shown on the kinase domain. (**B**) and (**C**) representation of the strategies used to produce full length and kinase domain recombinant PKCα. The kinase domain is cleaved with the TEV protease at the level of a TEV cleavage site (introduced at the residue 321) following expression of the full length protein. (**D**) activation loop phosphorylation (pT497) of full length WT-PKCα and D463H and D463N mutants. The addition of PDK1 kinase domain to increase phosphorylation efficiency of the turn motif, did not show further improvement of the phosphorylation status (the same is seen with the Turn motif and the HM in E). (**E**) Turn motif (pT638 (TM)) and hydrophobic motif (pS657 (HM)) phosphorylation of full length WT-PKCα. The phosphorylation was detected by western blot and the expression of the protein visualized by Coomassie blue staining.

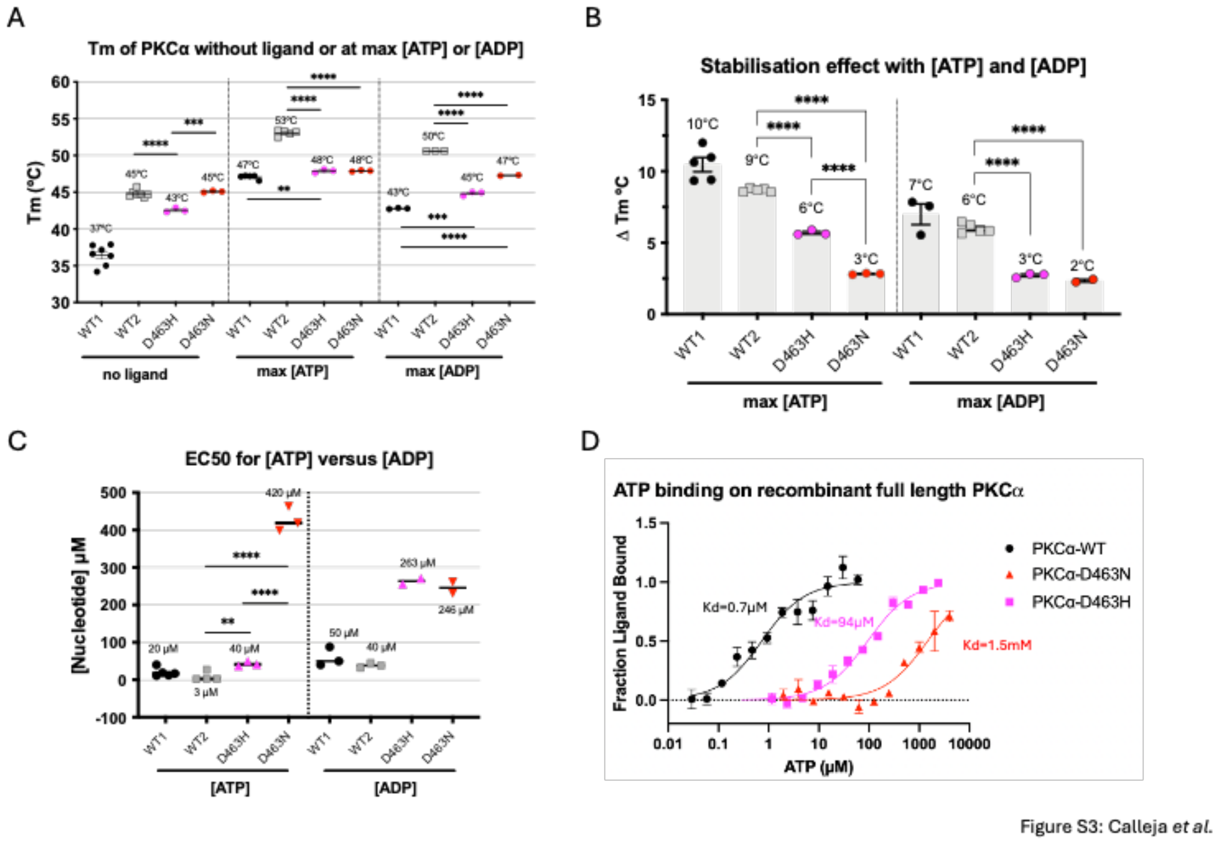


**Fig. S3. Thermal shift assay and microscale thermophoresis.** Recombinant WT-PKCα and mutants were mixed with ATP or ADP in presence of a fluorescent dye. The gradual increase in temperature triggers the denaturation of the recombinant protein and allows its labelling with the dyes. (**A**) melting temperatures (Tm) determined by the fitting the melting curves of recombinant WT-PKCα and mutants prior to or upon maximum ATP or ADP loading. The WT-PKCα has a biphasic denaturation curve (WT1 and WT2) while the two mutants show a monophasic curve. (**B**) The variation of the melting temperature (Tm) prior to (Apo) or in presence of nucleotides (ATP or ADP) allows the calculation of the ΔTm. A positive ΔTm represents an increase in the stability of the recombinant proteins obtained by the binding of the nucleotide. An unpaired t-test was used to determine the significance of the different changes. (**C**) Apparent affinities (EC50) calculated from the increase in Tm, of WT-PKCα versus D463H and D463N mutants for ATP or ADP. In all the TSA experiments the significance was assessed using an unpaired t-test P<0.01 (**), P<0.001(***), P<0.0001 (****) (each dot represents an independent experiment). (**D**) ATP binding affinities (Kd) were determined on full-length recombinant PKCαWT and mutants by microscale thermophoresis (n=3).

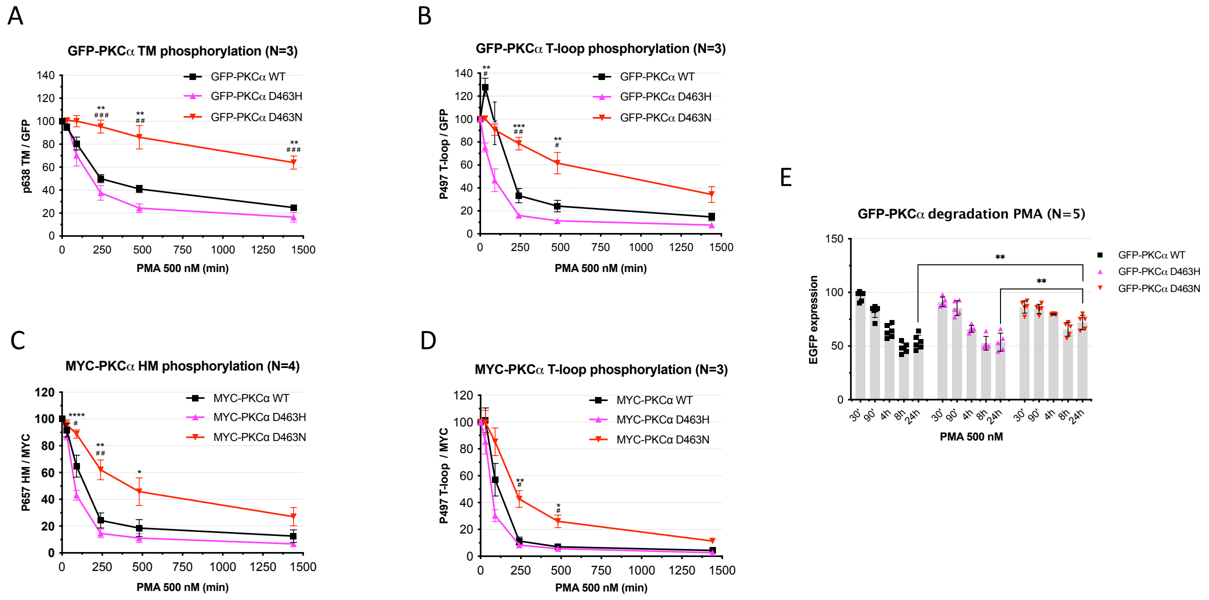


**Fig. S4. Similar regulation of WT-PKCα and D463H mutant constructs upon PMA treatment.** (**A-D**), GFP-PKCα (graphs A and B) and Myc-PKCα (graphs C and D) WT and mutants were transiently expressed in U87MG cells. The cells were treated with 500 nM PMA up to 24 hours. The decrease of the T-loop, hydrophobic motif or Turn motif phosphorylation is visible for WT-PKCα and mutant constructs. (**A**) GFP-PKCα WT and mutants decrease phosphorylation of the turn motif (TM) (n=3). Comparison of the TM phosphorylation of D463N vs D463H (p-values*) and D463N vs WT (p-values#). (**B**) GFP-PKCα WT and mutants decrease phosphorylation of the T-loop (n=3). The statistical comparison the T-loop phosphorylation was done for D463N vs D463H (p-values*) and D463N vs WT (p-values#). (**C**) Myc-PKCα WT and mutants decrease phosphorylation of the hydrophobic motif (HM) (n=4). Statistical analysis was done to compare the HM phosphorylation of D463N vs D463H (p-values*) and D463N vs WT (p-values#). (**D**) Myc-PKCα WT and mutants decrease phosphorylation of the T-loop (n=3). Statistical analysis to compare the T-loop phosphorylation of D463N vs D463H (p-values*) and D463N vs WT (p-value#). (**E**) shows the increased stability of the D463N mutant compared to WT and D463H upon prolonged PMA treatment (n=5). In all experiments the confidence was assessed using an unpaired t-test P<0.05 (*), P<0.01 (**), P<0.001(***), P<0.0001 (****).

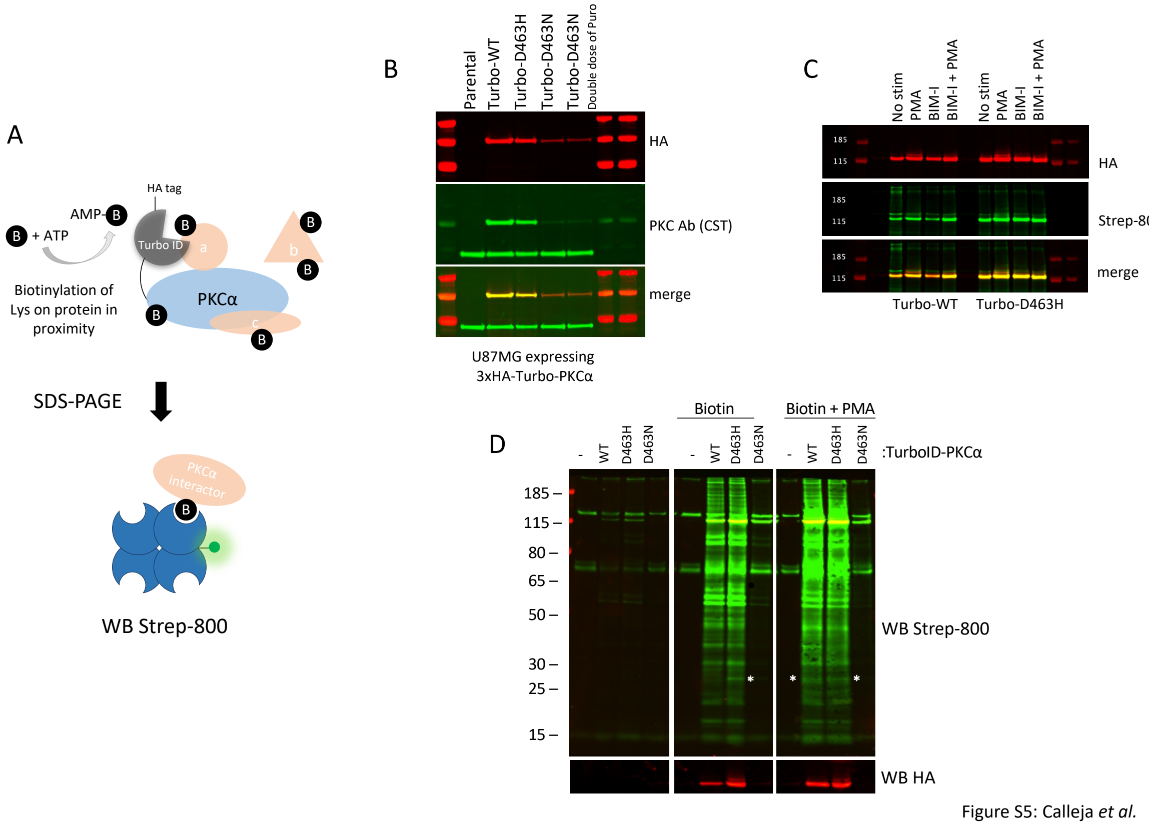


Figure S5: Calleja *et al.*

**Fig. S5. TurboID-PKCα biotinylation of PKCα and mutants.**

(**A**) Schematic principle of TurboID-tagged PKCα biotinylation (black round labelled with 'B') of associated proteins (light pink) and PKCα itself and recognition of the biotinylated proteins with streptavidin labelled with IRDye800. (**B**) Stable expression of 3xHA-TurboID-PKCα WT and mutants in U87MG cells detected by western blot with anti HA or anti PKCα antibodies. The expression of endogenous PKCα (lower band in WB PKCα) is indicative of equal amount of cell lysate loaded. The expression of D463N mutant is highly reduced compared to WT and D463H mutants and cannot be increased by increased selection pressure. (**C**) Western blot anti HA or Streptavidin following neutravidin pull down of 3xHA-TurboID-PKCα from U87MG stable cell lines. The expression (red) and biotinylation status (green) of 3xHA-TurboID-PKCα WT and mutants is shown upon treatment with PMA or BIM-I. (**D**) Western blot anti HA or fluorescently labelled Streptavidin following neutravidin pull down of 3xHA-TurboID-PKCα from U87MG stable cell lines. The neutravidin pull down of biotinylated proteins from cells stably expressing 3xHA-TurboID-PKCα WT and mutant D463H is compared to the pull down of untransfected parental U87MG cells prior or upon PMA treatment. The expression of PKCα is shown by western blot anti HA. The white stars indicate the protein differently pulled down in D463H conditions compared to WT.

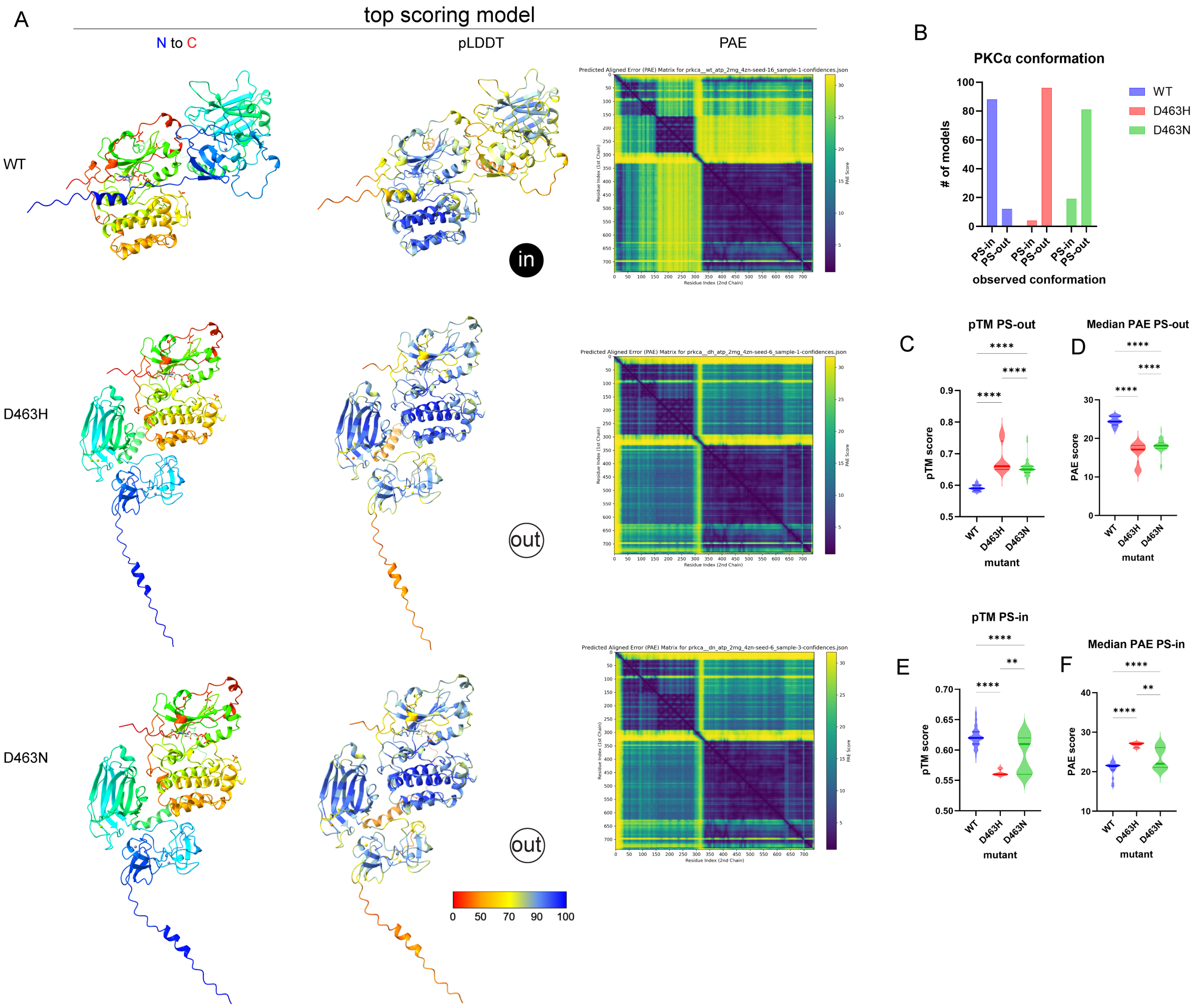


**Fig. S6. Calleja et al**

**Fig. S6. Structure predictions for WT and mutant forms of PKCα.** (A) Top scoring AlphaFold3 models for PKCα WT, D463H and D463N mutants, models shown in a rainbow colour scheme on the left and according to pLDDT score to the right. The PAE matrices are shown next to the models. the in/out denotation represents whether the model presents in a PS-in or PS-out conformation (see text). (B) Model classification per conformation (PS-in or PS-out). (C) pTM scores of models observed in a PS-out conformation for WT and mutant PKCαpredictions. (D) Median PAE scores for models observed in a PS-out conformation for WT and mutant PKCα predictions. (E) pTM scores of models observed in a PS-in conformation for WT and mutant PKCα predictions. (F) Median PAE scores for models observed in a PS-in conformation for WT and mutant PKCα predictions. Data in C-F are presented as violin plots and were analysed using one-way ANOVA followed by Tukey's multiple comparisons test. Statistical significance is indicated by ** $p \leq 0.01$, **** $p \leq 0.0001$.

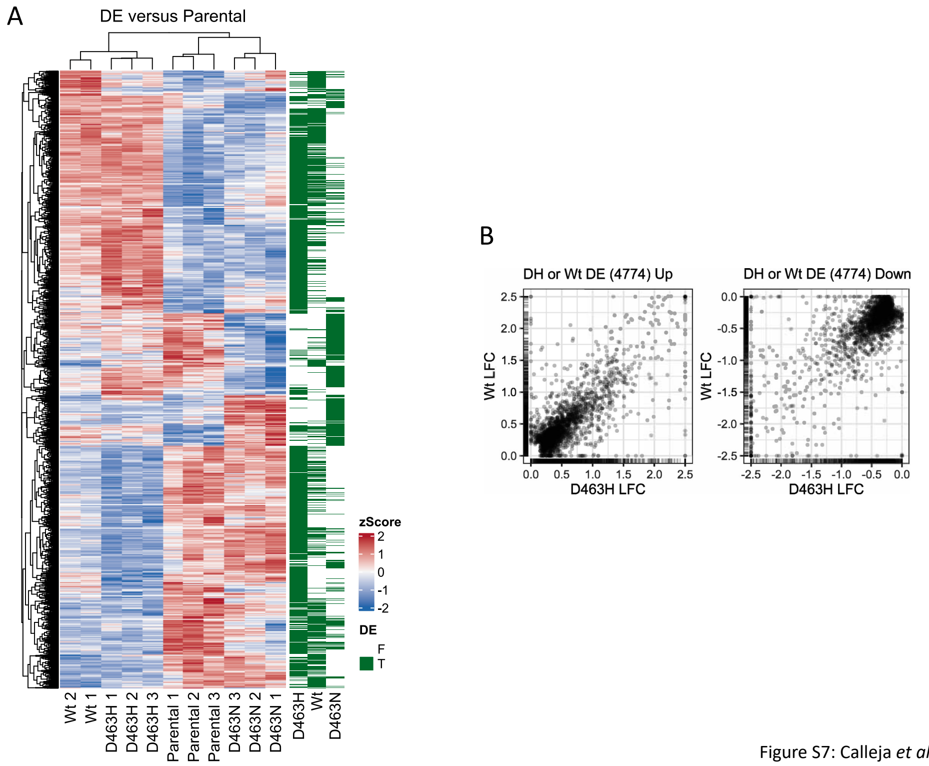


Figure S7: Calleja *et al.*

**Fig. S7. PKCα D463H is a gain-of-function mutant.** (**A**) Heatmap of genes differentially expressed with overexpression of Myc-tagged WT-PKCα, D463H and D463N compared to parental cells. Tiles are coloured by per-gene z-scores across all samples. Right heatmap annotation identifies significant differential gene expression for each group compared to parental. (**B**) Scatter plots describing the relationship in log fold change in gene expression between WT and D463H for genes differentially expressed in PKCα WT or D463H stable U87MG cells when compared with parental cell lines.

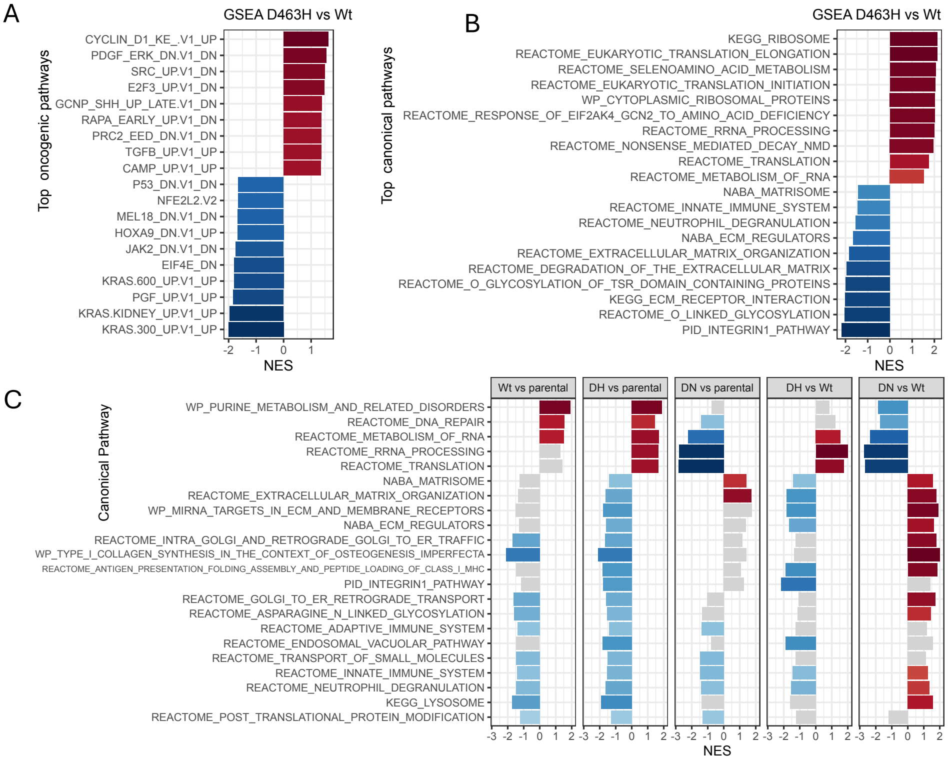


**Fig. S8. GSEA analysis of differential oncogenic and canonical pathways between WT-PKCα and D463H. (A-C)** Normalised enrichment scores from gene set enrichment analysis for the oncogenic pathways (A), canonical pathways (B, C) collections obtained from the molecular signatures database. Ranks were calculated based on the Wald statistic from differential expression results between D463H and WT-PKCα U87MG cells in A and B and in all differential expression comparisons in C. The top and bottom ten most significantly enriched pathways from the collections are shown in A and B. Gene sets with two or more agreement between the WT-PKCα versus parental, D463H versus parental and D463H versus WT-PKCα samples are shown in C.

#### Supplementary Tables

**Table S1. *Prkca*-D463H mosaic mice obtained from the first round of CRISPR/Cas9 genome editing.** From the first round of CRISPR in C57BL/6J mice, the mosaic male IRCP4.1c (in red) with 22% penetrance of the mutation D463H, survived and was used for timed mating.

| Mosaic ID | M/F | PRKCA mutation | Guide | Cell stage | Miseq results | Fate |
|---|---|---|---|---|---|---|
| IRCP1.1a | M | -- | 1 | 2C | Wildtype (>90%) | culled surplus |
| IRCP1.1b | M | -- | 1 | 2C | Wildtype (>90%) | culled surplus |
| IRCP1.1c | M | -- | 1 | 2C | Wildtype (>90%) | culled surplus |
| IRCP1.1d | F | -- | 1 | 2C | Wildtype (>90%) | culled surplus |
| IRCP3.1a | M | UD4 | 2 | 1C | Wildtype (71%); 1bp del (15%); 1bp ins (14%) | culled surplus |
| IRCP3.1b | M | UD4 | 2 | 1C | 65bp del (60%); PAM mut (40%) | culled sick |
| IRCP3.1c | M | -- | 2 | 1C | Indels | culled surplus |
| IRCP3.1d | M | +- | 2 | 1C | Desired mut (65%); 2bp deletion (25%), 4bp del (14%) | culled hydrocephalus |
| IRCP3.1e | F | ++ | 2 | 1C | Desired mut (22%); WT (43%), 108bp del (35%) | culled hydrocephalus |
| IRCP3.1f | F | ++ | 2 | 1C | Desired mut (>90%) | culled hydrocephalus |
| IRCP3.1g | F | +- | 2 | 1C | Desired mut (50%); WT (50%) | culled hydrocephalus |
| IRCP3.1h | F | ++ | 2 | 1C | 65bp del (67%); PAM mut (33%), | culled surplus |
| **IRCP3.1i** | **F** | **+-** | **2** | **1C** | **Desired mut (23%); 1bp ins (40%); 2bp ins (40%)** | **culled 22 w broken leg** |
| IRCP4.1a | M | +- | 2 | 1C | Desired mut (11%); 171bp del (60%); PAM mut (16%); WT (13%) | culled surplus |
| IRCP4.1b | M | -- | 2 | 1C | low read count, but only indels detected | culled sick |
| IRCP4.1c | M | UD4 | 2 | 1C | Desired mut (22%); WT (78%) | alive for 57 weeks |
| IRCP4.1d | F | ++ | 2 | 1C | PAM positive (75%); 4bp ins (25%) | found dead |
| **IRCP4.1e** | **F** | **+-** | **2** | **1C** | **Desired mut (26%); WT (29%); 1bp ins (25%);7bp del (19%)** | **culled 30 weeks emaciated** |
| IRCP4.1f | F | -- | 2 | 1C | WT (70%); 1bp ins (30%) | culled surplus |

**Table S2. *Prkca* D463H Het mice obtained from IRCP4.1c mosaic mice by timed mating.** From the timed mating of the male mosaic mouse IRCP4.1c (table S1) with C57BL/6J females 7 heterozygous mice were identified out of 46 pups. All 7 het mice displayed an adverse phenotype and had to be culled at 3 weeks.

| Founder ID cross | Litter name | Hets/Total pups | Phenotype |
|---|---|---|---|
| **IRCP4.1c** X C57BL/6J | PPBR5.1 | 0/8 | - |
| | PPBR6.1 | 1/7 | Hunchback and smaller |
| | PPBR7.2 | 2/7 | Hunchback and smaller |
| | PPBR8.2 | 0/8 | - |
| | PPBR9.1 | 1/8 | Hunchback and smaller |
| | PPBR10.1 | 3/8 | Hunchback and smaller |

**Table S3. *Prkca* D463H Het mice obtained from IRCP4.1c mosaic mice by IVF of CD-1.** CD-1 female mice were used for IVF with the mosaic male IRCP4.1c (table S1). Despite greater genetic diversity of the CD-1 background the 3 born heterozygotes from 4 litters displayed a deleterious phenotype. Only one D463H het male (PPBY4.1c) survived up to 11 weeks.

| Founder ID IVF | Litter name | Hets/Total pups | Het ID | Phenotype |
|---|---|---|---|---|
| **IRCP4.1c** on CD-1 | PPBY1.1 | 1/9 | PPBY1.1e (2weeks) | Hindlimb deformation |
| | PPBY1.1 | 0/11 | - | - |
| | PPBY4.1 | 1/8 | **PPBY4.1c (11 weeks)** | Hindlimb deformation |
| | PPBY5.1 | 1/5 | PPBY5.1a (2 weeks) | Hindlimb deformation |

**Table S4. *Prkca* D463H cross for Het mice with PPBY4.1c in trio with CD-1.** The D463H Het male PPBY4.1c was placed in trio with CD-1 female mice and produced 4 hets who did not survive later than 9 weeks, all displaying a harmful phenotype of hindlimb deformation.

| Het ID: het D463H x in trio with CD-1 | Litter name | Hets/Total pups | Phenotype |
|---|---|---|---|
| **PPBY4.1c** X 2 CD-1 | PPBY6 | 4/12 (9 weeks) | Hindlimb deformation |
| | PPBY7 | 0/0 | |

**Table S5. *Prkca* D463H Het mice obtained from 2 mosaic mice from second round of CRISPR/Cas9 genome editing.** New mosaic mice were obtained from a second round of CRISPR on a C57BL/6J background or on an F1(B6 x CBA) background (IRCP8.1e and IRCP20.1b respectively). Both mosaic males were crossed with two CD-1 females. The 21 *Prkca* D463H heterozygotes produced from both male's backgrounds were sickly and displayed a deformation of the hindlimbs.

| Founder ID cross in trio with CD-1 | Litter name | Hets/Total pups | Age of hets at cull | Phenotype |
|---|---|---|---|---|
| **IRCP8.1e** ♂ (12% mosaic ) X 2 CD-1 | PPCE1.1 | 8/13 | 3 weeks | Hindlimb deformation |
| | PPCE2.1 | 4/12 | 2 and 8 weeks | Hindlimb deformation |
| | PPCE2.2 | 3/14 | 3 weeks | Hindlimb deformation |
| **IRCP20.1b** ♂ (18% mosaic) X 2 CD-1 | PPCE7.1 | 2/13 | 3 and 9 weeks | Hindlimb deformation |
| | PPCE8.1 | 4/10 | 11 weeks | Hindlimb deformation |

**Table S6. *Prkca*-D463N control mosaic mice obtained from CRISPR/Cas9 genome editing.** The same guide RNAs used for the D463H mutation but with a modified repair template was used to produce mosaic mice harbouring the control mutation D463N. 16 mosaic C57BL/6J mice were obtained with the mutation D463N, and 3 of them (IRCY3.1h, IRCY6.1b and IRCY6.1L) were bred for germline transmission.

| Founder mosaic ID | M/F | Guide | PRKCA mutation D>N | Miseq summary | Fate |
|---|---|---|---|---|---|
| IRCY3.1b | M | 1 | +- | Deletion alleles, minor allele with pm** HA duplicated | Culled |
| IRCY3.1d | F | 1 | UD4 | Incorrect insertion alleles, minor allele with pm | Culled |
| IRCY3.1e | F | 1 | UD4 | Hom for 122bp del, minor allele with pm | Culled |
| IRCY3.1g | F | 1 | ++ | Hom for pm, sm absent, minor indel allele | Culled |
| **IRCY3.1h** | **F** | **1** | **UD4** | **Het for pm, sm absent, wt allele with indel** | **Bred** |
| IRCY6.1a | M | 2 | +- | Bias to wt allele but allele with pm and sm present | Culled |
| **IRCY6.1b** | **M** | **2** | **UD4** | **Perfect het, wt contains indel** | **Bred** |
| IRCY6.1c | M | 2 | +- | Perfect het, wt contains indel | Stored |
| IRCY6.1d | M | 2 | UD4 | Perfect hom | Stored |
| IRCY6.1f | M | 2 | ++ | Hom, no sm | Stored |
| IRCY6.1g | F | 2 | UD4 | Low reads, likely perfect het with sm | Culled |
| IRCY6.1i | F | 2 | ++ | Het, no sm | Stored |
| IRCY6.1k | F | 2 | ++ | Het + sm, further pm allele contains a duplication | Stored |
| **IRCY6.1l** | **F** | **2** | **UD4** | **Perfect het with sm, wt contains indel** | **Bred** |
| IRCY6.1m | F | 2 | UD4 | Perfect het with sm, wt contains indel | Stored |
| IRCY9.1a | M | 2 | UD4 | Probable hom, no sm | Stored |

**Table S7. *Prkca* D463N control Het mice obtained from three mosaic mice from CRISPR/Cas9 genome editing.** Three mosaic D463N mice (IRCY3.1h, 6.1b and 6.1l) were crossed with C57BL/6J mice to obtain 15 *Prkca* D463N heterozygous mice. All the D463N het mice were asymptomatic.

| Founder mosaic ID cross | Litter name | Hets/Total pups | Phenotype |
|---|---|---|---|
| IRCY3.1h ♀ X C57BL/6J | IRCY13.1 | 4/9 | none |
| IRCY6.1b ♂ X C57BL/6J | IRCY11.1 | 3/8 | none |
| | IRCY11.2 | 3/6 | none |
| IRCY6.1l ♂ X C57BL/6J | IRCY12.1 | 5/7 | none |

**Table S8.** ***Prkca*** **D463N control second round of cross for Het mice.** *Prkca* D463N het offspring from the litters IRCY11, IRCY12 or IRCY13 (see table S7) were crossed with either C57BL/6J, F1(B6 x CBA) or CD-1 mice to obtain a second round of *Prkca* D463N heterozygotes (litters PPBA, PPCB and PPBZ). All the het showed a normal phenotype and good health.

| Het ID: het D463N x WT breeders | Litter name | Hets/Total pups | Phenotype |
|---|---|---|---|
| IRCY11.1g ♀ X C57BL/6J | PPBZ2.1 | 4/8 | none |
| IRCY11.1g ♀ X C57BL/6J | PPBZ2.3 | 3/8 | none |
| IRCY11.1h ♀ X C57BL/6J | PPBZ3.1 | 1/2 | none |
| IRCY11.2b ♂ X F1(B6 x CBA) | PPBZ7.1 | 6/8 | none |
| IRCY12.1g ♀ X C57BL/6J | PPCA1.1 | 3/3 | none |
| IRCY12.1b ♂ X CD-1 | PPCA4.1 | 9/15 | none |
| IRCY12.1c ♂ X C57BL/6J | PPCA11.1 | 2/8 | none |
| IRCY13.1d ♀ X C57BL/6J | PPCB1.1 | 2/6 | none |

**Table S9.** ***Prkca*** **D463N control cross for Homozygous mice.** *Prkca* D463N het mice from 3 different litters (see table S7) were crossed. From two different crosses, 12 homozygotes, 14 heterozygotes and 6 WT pups were generated (litters PPBZ8, PPCA12 and PPCB12). None of the mice displayed any phenotype.

| Het ID: het D463N x het D463N | Litter name | Hom/Total pups | Hets/Total pups | WT/Total pups | Phenotype |
|---|---|---|---|---|---|
| IRCY11.2a (♂) X IRCY13.1d (♀) | PPBZ8.1 | 2/9 | 7/9 | 0/9 | none |
| | PPBZ8.3 | 4/9 | 3/9 | 2/9 | none |
| IRCY12.1b (♂) X IRCY11.1h (♀) | PPCA12.1 | 3/5 | 0/5 | 2/5 | none |
| | PPCB12.2 | 2/4 | 2/4 | 0/4 | none |
| | PPCB12.3 | 1/5 | 2/5 | 2/5 | none |

**Table S10. Guide RNAs for knock-in studies.**

| | |
|---|---|
| Guide RNA 1: | CCCTTGCAGGGATCTGAAGC |
| Guide RNA 2: | CACTCACCTGCTCCCTTGCA |
| D463H ssODN | TGCAAGCAAATCAAGTCATGTCGTGGCTCGGCCTCTGACACTCACCTGCTCCCTTGCAGGcATCT GAAGCTcGACAATGTCATGCTGGACTCAGAAGGGCACATCAAAATCGCCGACTTTG |
| D463N ssODN | TGCAAGCAAATCAAGTCATGTCGTGGCTCGGCCTCTGACACTCACCTGCTCCCTTGCAGaATCTG AAGCTcGACAATGTCATGCTGGACTCAGAAGGGCACATCAAAATCGCCGACTTTG |

**Table S11. Oligonucleotide sequences for construct cloning.**

| **Mammalian cell expression** | **Oligo sequences used for PCR amplification** |
|---|---|
| Myc-PKCα_sense | CAGTGTGCTGGAATTCGCCGCCACCATGGAACAAAAACTCATCTCAGAAGAGGATCT GGCTGACGTTTTCCCGG |
| Myc-PKCα_anti | GATATCTGCAGAATTCTCATACTGCACTCTGTAAGATGGGG |
| Myc-PKCα_for | GCTCGAGCCCGAATTCGCCGCCACCATGGAACAAAAAC |
| Myc-PKCα_rev | GAGATCTCCCGAATTCTCATACTGACTCTGTAAGATGGGG |
| 3xHA-TurboID_for | TCCGGGCTCGAGCCCGAATTCGCCGCCACCATGTACCCG |
| 3xHA-TurboID_rev | CCTCCACCTCCACCCTTTTCGGCAGACCGCAGACTG |
| 6xGly-PKCα_for | GAAAAGGGTGGAGGTGGAGGAGGCGCTGACGTTTTCCCGGGCAAC |
| 6xGly-PKCα_rev | GATGGGAGATCTCCCGAATTCTCATACTGCACTCTGTAAGATGGGG |
| **Insect cell expression** | **Oligo sequences used for PCR amplification** |
| GST-3C PKCα_for | CCCTAAGCTTGGATCCTATGGCTGACGTTTTCCCGGGCAACGAC |
| GST-3C PKCα_rev | GGCCGCCCGGGAATTCTCATACTGCACTCTGTAAGATGGGGTG |
| PKCα Reg_for | CCCTAAGCTTGGATCCTATGGCTGACGTTTTCCCGGGCAACGAC |
| PKCα Reg_rev | TTCAGAGGGACTGATGACTTTGTTGCCAGCAGGGCCAAG |
| TEV321_sense | ATCAGTCCCTCTGAAAACCTCTACTTCCAGGGCGACAGGAAACAACCT |
| TEV321_anti | AGGTTGTTTCCTGTCGCCCTGGAAGTAGAGGTTTTCAGAGGGACTGAT |
| PKCα_Kin_for | GACAGGAAACAACCTTCCAACAACCTTGACCGAGTG |
| PKCα Kin_rev | GGCCGCCCGGGAATTCTCATACTGCACTCTGTAAGATGGGGTG |

**Table S12. Interactors PKCα WT, PKCα-D463H, or PKCα-D463N.** Proteins co-immunoprecipitated with MYC-tagged PKCαWT and mutants D463H and D463N from U87MG cells were analysed by label free quantitative mass spectrometry**.** All binding partners pulled down by WT or either D463 mutant, which are not found in any the control precipitates, are presented as the LFQ intensities; data are also presented as a clustered heat map (Figure 4D).

| my_id | id | gene | control_1 | control_2 | control_3 | WT_1 | WT_2 | WT_3 | D463H_1 | D463H_2 | D463H_3 | D463N_1 | D463N_2 | D463N_3 | control_ave | WT_average | D463H_ave | D463N_ave | group |
|---|---|---|---|---|---|---|---|---|---|---|---|---|---|---|---|---|---|---|---|
| 3 | B7ZAB0;B7Z4X6;A0A02 | SERPINE1 | 0 | 0 | 0 | 435320 | 930290 | 730560 | 475580 | 540610 | 577030 | 0 | 0 | 466530 | 0 | 698723.3 | 531073.3 | 155510 | all_PKCa_(66) |
| 4 | B3KNZ4;B4E2Z6;B4DD | EIF3C;EIF3 | 0 | 0 | 0 | 409770 | 421380 | 421720 | 435330 | 0 | 415860 | 407070 | 578450 | 433890 | 0 | 417623.3 | 283730 | 473136.7 | all_PKCa_(66) |
| 31 | Q5JR06;Q5JR07;E9PN1 | RHOC;RHO | 0 | 0 | 0 | 342790 | 575410 | 0 | 309220 | 338330 | 288900 | 362400 | 0 | 0 | 0 | 306066.7 | 312150 | 120800 | all_PKCa_(66) |
| 43 | A0A024R1K7;Q04917 | YWHAH | 0 | 0 | 0 | 614800 | 828450 | 588150 | 0 | 743790 | 567310 | 0 | 0 | 773630 | 0 | 677133.3 | 437033.3 | 257876.7 | all_PKCa_(66) |
| 52 | Q6ZME7;Q658U4;A0A( | DKFZp666[ | 0 | 0 | 0 | 728290 | 839260 | 645750 | 1127100 | 914850 | 670280 | 635560 | 662860 | 595720 | 0 | 737766.7 | 904076.7 | 631380 | all_PKCa_(66) |
| 53 | A0A0S2Z487;A0A024R | JUP | 0 | 0 | 0 | 432510 | 482550 | 577010 | 744010 | 873160 | 928850 | 509680 | 913570 | 1893300 | 0 | 497356.7 | 848673.3 | 1105517 | all_PKCa_(66) |
| 68 | A0A024R326;Q6IPI1;P | RPL29 | 0 | 0 | 0 | 4201700 | 7452200 | 3521600 | 7676200 | 5053900 | 2726500 | 4282100 | 3226400 | 3141500 | 0 | 5058500 | 5152200 | 3550000 | all_PKCa_(66) |
| 83 | H0Y4K8;Q9UQS6;O956 | FN1;DKFZp | 0 | 0 | 0 | 0 | 0 | 254820 | 359450 | 0 | 0 | 249950 | 214970 | 253570 | 0 | 84940 | 119816.7 | 239496.7 | all_PKCa_(66) |
| 101 | B2RAQ8;A0A024R5F4; | CPT1A | 0 | 0 | 0 | 521200 | 267740 | 434490 | 548110 | 775770 | 486870 | 0 | 1920400 | 0 | 0 | 407810 | 603583.3 | 640133.3 | all_PKCa_(66) |
| 120 | B2RDX0;A8K7Z9;A0A0 | SHCBP1 | 0 | 0 | 0 | 515200 | 697970 | 597140 | 369060 | 468460 | 405940 | 695580 | 322350 | 358340 | 0 | 603436.7 | 414486.7 | 458756.7 | all_PKCa_(66) |
| 124 | B4DFL1;E9PEX6;B4DM | DLD | 0 | 0 | 0 | 156480 | 282820 | 253130 | 225900 | 268210 | 338050 | 189190 | 114300 | 297710 | 0 | 230810 | 277386.7 | 200400 | all_PKCa_(66) |
| 158 | A0A024R920;Q59FG1; | CACNA1E | 0 | 0 | 0 | 3052700 | 1645900 | 2106600 | 1151300 | 513860 | 568960 | 1236300 | 607170 | 343740 | 0 | 2268400 | 744706.7 | 729070 | all_PKCa_(66) |
| 218 | A0A024RDG5;Q53EU6 | HMFN0839 | 0 | 0 | 0 | 0 | 165270 | 0 | 260770 | 437030 | 324800 | 0 | 173910 | 94200 | 0 | 55090 | 340866.7 | 89370 | all_PKCa_(66) |
| 249 | A0A087WV29;Q9H0A0 | NAT10 | 0 | 0 | 0 | 173830 | 0 | 206980 | 0 | 0 | 243750 | 245120 | 0 | 0 | 0 | 126936.7 | 81250 | 81706.67 | all_PKCa_(66) |
| 292 | B2R7P6;A0A0A0MSW4 | PITPNB | 0 | 0 | 0 | 1364800 | 1325000 | 1567600 | 0 | 1103200 | 356820 | 0 | 1287800 | 1833200 | 0 | 1419133 | 486673.3 | 1040333 | all_PKCa_(66) |
| 295 | A0A140VKB1;A0A0A0N | CUL2 | 0 | 0 | 0 | 0 | 284390 | 0 | 299700 | 409480 | 417700 | 0 | 0 | 323280 | 0 | 94796.67 | 375626.7 | 107760 | all_PKCa_(66) |
| 298 | F8WDD7;Q9H7Z5;H7C | ARPC4;ARF | 0 | 0 | 0 | 185300 | 427540 | 388890 | 355710 | 445720 | 516340 | 182710 | 0 | 0 | 0 | 333910 | 439256.7 | 60903.33 | all_PKCa_(66) |
| 301 | I3L1U7;I3L0X9;I3L3T0 | PAM16;CO | 0 | 0 | 0 | 258410 | 319140 | 232290 | 0 | 305410 | 321550 | 182400 | 115000 | 137530 | 0 | 269946.7 | 208986.7 | 144976.7 | all_PKCa_(66) |
| 316 | H0YA55;B4DPR2;F6KP | ALB | 0 | 0 | 0 | 1884400 | 2367100 | 2300200 | 2538200 | 2809700 | 2266700 | 2347800 | 2483900 | 2314000 | 0 | 2183900 | 2538200 | 2381900 | all_PKCa_(66) |
| 322 | H7BY36;A0A0D9SFL3;I | EWSR1 | 0 | 0 | 0 | 632200 | 547810 | 578270 | 0 | 386710 | 505320 | 495650 | 499550 | 554290 | 0 | 586093.3 | 297343.3 | 516496.7 | all_PKCa_(66) |
| 323 | A0A0F6MY87;Q8IVF6;A | ANKRD18A | 0 | 0 | 0 | 4272300 | 3792000 | 5358200 | 2609300 | 2810000 | 2680200 | 3062600 | 3201500 | 0 | 0 | 4474167 | 2699833 | 2088033 | all_PKCa_(66) |
| 347 | K7ESM3;K7EKQ5;K7EF | LGALS3BP | 0 | 0 | 0 | 106520 | 105550 | 107280 | 0 | 110940 | 0 | 0 | 96526 | 0 | 0 | 106450 | 36980 | 32175.33 | all_PKCa_(66) |
| 429 | I0J7B9;J3KR33;A0JLT2 | MED19AS;I | 0 | 0 | 0 | 419610 | 384470 | 177440 | 286140 | 195770 | 192570 | 0 | 126630 | 0 | 0 | 327173.3 | 224826.7 | 42210 | all_PKCa_(66) |
| 440 | Q4VB24;B2R984;A3R0 | HIST1H1E; | 0 | 0 | 0 | 909470 | 1007300 | 1015900 | 0 | 1196400 | 864840 | 0 | 1298500 | 1182100 | 0 | 977556.7 | 687080 | 826866.7 | all_PKCa_(66) |
| 465 | B5B2S5;B5B2S3;B5B2 | NFATC3 | 0 | 0 | 0 | 210840 | 0 | 0 | 0 | 226880 | 308680 | 187890 | 184600 | 219130 | 0 | 70280 | 178520 | 197206.7 | all_PKCa_(66) |
| 508 | A8K9U0;Q9BWF3;E9PE | RBM4 | 0 | 0 | 0 | 0 | 0 | 361550 | 531450 | 373750 | 366900 | 494780 | 401860 | 587730 | 0 | 120516.7 | 424033.3 | 494790 | all_PKCa_(66) |
| 536 | B2R5T1;B7Z3W6;P051 | PRKCG | 0 | 0 | 0 | 375040 | 534840 | 566580 | 610440 | 627910 | 656370 | 280290 | 215790 | 308030 | 0 | 492153.3 | 631573.3 | 268036.7 | all_PKCa_(66) |
| 539 | B4DSE2;B4E0C7;Q53H | TPP1 | 0 | 0 | 0 | 0 | 235960 | 186740 | 165460 | 198530 | 0 | 121480 | 0 | 0 | 0 | 140900 | 121330 | 40493.33 | all_PKCa_(66) |
| 545 | B2R6S9;P30533;A8K8 | LRPAP1 | 0 | 0 | 0 | 0 | 0 | 1339200 | 1186600 | 1324200 | 1142800 | 1112300 | 1136200 | 1084500 | 0 | 446400 | 1217867 | 1111000 | all_PKCa_(66) |
| 573 | B3KMP6;D3DSW3;O94 | PROSC | 0 | 0 | 0 | 435090 | 372670 | 290140 | 373370 | 321920 | 263380 | 181080 | 0 | 154040 | 0 | 365966.7 | 319556.7 | 111706.7 | all_PKCa_(66) |
| 585 | Q5H9A7;H0Y789;Q6FG | TIMP1 | 0 | 0 | 0 | 487880 | 580200 | 870200 | 1352900 | 0 | 929460 | 0 | 0 | 433880 | 0 | 646093.3 | 760786.7 | 144626.7 | all_PKCa_(66) |
| 628 | B4DRW1;B4DRR7;B4D | KRT4 | 0 | 0 | 0 | 3878300 | 1952500 | 3525000 | 4333900 | 4442500 | 7694000 | 0 | 0 | 2572200 | 0 | 3118600 | 5490133 | 857400 | all_PKCa_(66) |
| 649 | B4DQ50;Q08188 | TGM3 | 0 | 0 | 0 | 0 | 328360 | 408250 | 227100 | 0 | 419990 | 0 | 691870 | 0 | 0 | 245536.7 | 215696.7 | 230623.3 | all_PKCa_(66) |
| 658 | B4DUG4;K7ELG9;Q3M | LSM12 | 0 | 0 | 0 | 706520 | 636440 | 599110 | 718980 | 662290 | 619620 | 621960 | 734180 | 334440 | 0 | 647356.7 | 666963.3 | 563526.7 | all_PKCa_(66) |
| 670 | H0YEC5;H0YD77;H0YC | EIF4G2;AA( | 0 | 0 | 0 | 384130 | 410810 | 401730 | 297490 | 0 | 311740 | 483360 | 665620 | 481420 | 0 | 398890 | 203076.7 | 543466.7 | all_PKCa_(66) |
| 718 | G9I2K8;C7FDR3 | HLA-B | 0 | 0 | 0 | 178780 | 300040 | 286350 | 332680 | 365480 | 310980 | 220020 | 223090 | 345330 | 0 | 255056.7 | 336380 | 262813.3 | all_PKCa_(66) |
| 720 | C9J712;P35080;C9J2N | PFN2 | 0 | 0 | 0 | 2040100 | 1557600 | 0 | 2399400 | 1684700 | 0 | 1694300 | 1772100 | 0 | 0 | 1199233 | 1361367 | 1155467 | all_PKCa_(66) |
| 723 | C9JA28;Q9UNL2 | SSR3 | 0 | 0 | 0 | 288460 | 306990 | 317740 | 234730 | 296080 | 230630 | 0 | 223080 | 185780 | 0 | 304396.7 | 253813.3 | 136286.7 | all_PKCa_(66) |
| 730 | C9JUP3;E9PHG3;Q5HY | SMYD1;DKI | 0 | 0 | 0 | 0 | 2626500 | 3631200 | 0 | 2543100 | 2934300 | 0 | 4682700 | 5083500 | 0 | 2085900 | 1825800 | 3255400 | all_PKCa_(66) |
| 757 | CON__Q3TTY5 | CON__Q3T | 0 | 0 | 0 | 6064700 | 5990900 | 5552800 | 0 | 2458000 | 2301100 | 5041800 | 5569900 | 5175400 | 0 | 5869467 | 1586367 | 5262367 | all_PKCa_(66) |
| 769 | Q6NX58;D3DPS3;Q3K( | MAP7D1;RI | 0 | 0 | 0 | 346550 | 297690 | 381470 | 547530 | 639580 | 405540 | 654630 | 0 | 407960 | 0 | 341903.3 | 530883.3 | 354196.7 | all_PKCa_(66) |
| 781 | D6R9I9;P61221 | ABCE1 | 0 | 0 | 0 | 0 | 84771 | 132240 | 92056 | 0 | 0 | 0 | 0 | 83291 | 0 | 72337 | 30685.33 | 27763.67 | all_PKCa_(66) |
| 783 | D6RD66;V9HWG7;Q53 | WDR1;HEL | 0 | 0 | 0 | 0 | 377810 | 352070 | 0 | 374710 | 453900 | 405630 | 477170 | 414620 | 0 | 243293.3 | 276203.3 | 432473.3 | all_PKCa_(66) |
| 841 | G3V2R9;V9HW32;Q8N | PTGR2;HEL | 0 | 0 | 0 | 352770 | 293230 | 0 | 623010 | 440080 | 444520 | 0 | 0 | 296810 | 0 | 215333.3 | 502536.7 | 98936.67 | all_PKCa_(66) |
| 901 | K7EJ44;P07737 | PFN1 | 0 | 0 | 0 | 312010 | 222990 | 265550 | 386150 | 339250 | 0 | 247530 | 331420 | 302220 | 0 | 266850 | 241800 | 293723.3 | all_PKCa_(66) |
| 939 | Q6FHJ5;O14828 | SCAMP3 | 0 | 0 | 0 | 824480 | 861150 | 0 | 0 | 0 | 487990 | 431540 | 238590 | 400910 | 0 | 561876.7 | 162663.3 | 357013.3 | all_PKCa_(66) |
| 950 | Q5F0I5;Q5F0I4;O6039 | GRIN3B | 0 | 0 | 0 | 435850 | 341070 | 372090 | 75736000 | 71828000 | 6853700 | 406410 | 0 | 0 | 0 | 383003.3 | 51472567 | 135470 | all_PKCa_(66) |
| 994 | Q8TBT6;P08574 | CYC1 | 0 | 0 | 0 | 948220 | 557130 | 842160 | 659270 | 817310 | 470850 | 175300 | 0 | 0 | 0 | 782503.3 | 649143.3 | 58433.33 | all_PKCa_(66) |
| 1048 | P27694;I3L4R8 | RPA1 | 0 | 0 | 0 | 0 | 0 | 169530 | 0 | 0 | 158730 | 101590 | 172050 | 0 | 0 | 56510 | 52910 | 91213.33 | all_PKCa_(66) |
| 1062 | P35606 | COPB2 | 0 | 0 | 0 | 410160 | 306680 | 318090 | 366380 | 276100 | 362430 | 366140 | 344640 | 335210 | 0 | 344976.7 | 334970 | 348663.3 | all_PKCa_(66) |
| 1069 | P40222 | TXLNA | 0 | 0 | 0 | 461580 | 380860 | 0 | 538560 | 355290 | 293700 | 407630 | 316350 | 427210 | 0 | 280813.3 | 395850 | 383730 | all_PKCa_(66) |
| 1100 | P54886 | ALDH18A1 | 0 | 0 | 0 | 406180 | 326680 | 0 | 0 | 410140 | 397460 | 452900 | 416120 | 0 | 0 | 244286.7 | 269200 | 289673.3 | all_PKCa_(66) |
| 1102 | Q53FD2;P57060 | RWDD2B | 0 | 0 | 0 | 376240 | 525510 | 498880 | 259880 | 301880 | 334440 | 202030 | 188970 | 218450 | 0 | 466876.7 | 298733.3 | 203150 | all_PKCa_(66) |
| 1180 | Q13518;Q9UG07;Q8W | DKFZp586I | 0 | 0 | 0 | 0 | 1983100 | 2007100 | 2103200 | 2253000 | 0 | 0 | 1655800 | 1718800 | 0 | 1330067 | 1452067 | 1124867 | all_PKCa_(66) |
| 1191 | Q14693;Q68CS2;A0A( | LPIN1;DKF | 0 | 0 | 0 | 346990 | 0 | 636210 | 604850 | 0 | 0 | 112080 | 70889 | 0 | 0 | 327733.3 | 201616.7 | 60989.67 | all_PKCa_(66) |
| 1194 | Q14964;C9JB90;C9JU1 | RAB39A;RA | 0 | 0 | 0 | 1303600 | 1370900 | 1298400 | 1228400 | 1259800 | 1166500 | 914380 | 1004000 | 1299700 | 0 | 1324300 | 1218233 | 1072693 | all_PKCa_(66) |
| 1206 | Q15654;Q68D94;F2ZC | TRIP6;DKF | 0 | 0 | 0 | 681220 | 536360 | 550830 | 734370 | 664150 | 597020 | 0 | 356990 | 0 | 0 | 589470 | 665180 | 118996.7 | all_PKCa_(66) |
| 1230 | Q58FF6 | HSP90AB4 | 0 | 0 | 0 | 811180 | 0 | 2059800 | 3165700 | 1313600 | 973320 | 1147600 | 2192500 | 1753000 | 0 | 956993.3 | 1817540 | 1697700 | all_PKCa_(66) |
| 1231 | Q58FF8 | HSP90AB2 | 0 | 0 | 0 | 20482000 | 20856000 | 16286000 | 21456000 | 23940000 | 21814000 | 8174200 | 6763800 | 7406600 | 0 | 19208000 | 22403333 | 7448200 | all_PKCa_(66) |
| 1263 | Q96G43;Q6PID8;C9JR | KLHDC10 | 0 | 0 | 0 | 1675400 | 1372300 | 1071800 | 1843000 | 1800000 | 1736200 | 1343200 | 1809600 | 1494200 | 0 | 1373167 | 1793067 | 1549000 | all_PKCa_(66) |
| 1332 | Q96QR8 | PURB | 0 | 0 | 0 | 0 | 0 | 152400 | 0 | 204380 | 0 | 143170 | 0 | 0 | 0 | 50800 | 68126.67 | 47723.33 | all_PKCa_(66) |
| 1351 | Q9H4B7 | TUBB1 | 0 | 0 | 0 | 7100600 | 8191600 | 3960400 | 2913000 | 3114600 | 2995100 | 7496600 | 6524600 | 7107600 | 0 | 6417533 | 3007567 | 7042933 | all_PKCa_(66) |
| 1374 | Q9UIW2;B4DE20;A0A( | PLXNA1 | 0 | 0 | 0 | 4738400 | 7076000 | 4818200 | 0 | 0 | 3303700 | 3859800 | 0 | 0 | 0 | 5544200 | 1101233 | 1286600 | all_PKCa_(66) |
| 1375 | Q9UJZ1;A0A087WYB4 | STOML2 | 0 | 0 | 0 | 200570 | 249080 | 271670 | 223680 | 0 | 275840 | 217160 | 0 | 0 | 0 | 240440 | 166506.7 | 72386.67 | all_PKCa_(66) |
| 1398 | REV__B3KVM6;REV__Q | REV__B3KV | 0 | 0 | 0 | 1352600 | 1302400 | 2594100 | 2647200 | 2769300 | 2020500 | 2561000 | 3535800 | 2056500 | 0 | 1749700 | 2479000 | 2717767 | all_PKCa_(66) |
| 1407 | REV__Q7Z2Z1 | REV__Q7Z2 | 0 | 0 | 0 | 5587900 | 10171000 | 9352000 | 2822200 | 2623200 | 2512000 | 3833700 | 1862200 | 3817400 | 0 | 8370300 | 2652467 | 3171100 | all_PKCa_(66) |
| 111 | Q5D0E5;G3V3Z8;A0A0 | NUMB | 0 | 0 | 0 | 0 | 0 | 0 | 0 | 0 | 469370 | 789480 | 0 | 0 | 0 | 0 | 156456.7 | 263160 | D463H_D463N_(14) |
| 159 | A0A024R978;Q9BZQ8 | C1orf24;FA | 0 | 0 | 0 | 0 | 0 | 0 | 187390 | 0 | 0 | 0 | 0 | 191210 | 0 | 0 | 62463.33 | 63736.67 | D463H_D463N_(14) |
| 173 | Q8N6F2;B4DKI9;Q6PK | MRPS27 | 0 | 0 | 0 | 0 | 0 | 0 | 320300 | 0 | 0 | 341800 | 343230 | 275430 | 0 | 0 | 106766.7 | 320153.3 | D463H_D463N_(14) |
| 179 | A0A024RAV4;A0A024F | CSDA;YBX3 | 0 | 0 | 0 | 0 | 0 | 0 | 0 | 0 | 561570 | 0 | 723040 | 965280 | 0 | 0 | 187190 | 562773.3 | D463H_D463N_(14) |
| 281 | B2RB99;B4DXL5;A0A0 | RTFDC1 | 0 | 0 | 0 | 0 | 0 | 0 | 0 | 403120 | 502640 | 0 | 183060 | 0 | 0 | 0 | 301920 | 61020 | D463H_D463N_(14) |
| 438 | B4DM79;Q5T985;CON | ITIH2 | 0 | 0 | 0 | 0 | 0 | 0 | 0 | 0 | 232320 | 161190 | 224740 | 198140 | 0 | 0 | 77440 | 194690 | D463H_D463N_(14) |
| 635 | B4DLA6 | B4DLA6 | 0 | 0 | 0 | 0 | 0 | 0 | 0 | 29351000 | 0 | 45198000 | 0 | 0 | 0 | 0 | 9783667 | 15066000 | D463H_D463N_(14) |
| 725 | C9JG08 | C9JG08 | 0 | 0 | 0 | 0 | 0 | 0 | 0 | 1576800 | 0 | 1183900 | 1653900 | 0 | 0 | 0 | 525600 | 945933.3 | D463H_D463N_(14) |
| 807 | E9PI41;Q9NPD3 | EXOSC4 | 0 | 0 | 0 | 0 | 0 | 0 | 381410 | 343560 | 0 | 295060 | 305810 | 420700 | 0 | 0 | 241656.7 | 340523.3 | D463H_D463N_(14) |
| 923 | M0QYE0;M0R0I3;Q9U( | SH3GL1;SH | 0 | 0 | 0 | 0 | 0 | 0 | 335300 | 0 | 329160 | 297870 | 257730 | 0 | 0 | 0 | 221486.7 | 185200 | D463H_D463N_(14) |
| 967 | O95714;A8KAQ8;A0A0 | HERC2 | 0 | 0 | 0 | 0 | 0 | 0 | 0 | 0 | 1553900 | 0 | 443930 | 0 | 0 | 0 | 517966.7 | 147976.7 | D463H_D463N_(14) |
| 1006 | P11388 | TOP2A | 0 | 0 | 0 | 0 | 0 | 0 | 0 | 56499 | 0 | 78859 | 90818 | 57138 | 0 | 0 | 18833 | 75605 | D463H_D463N_(14) |
| 1020 | P17252-mutantDN | PRKCA | 0 | 0 | 0 | 0 | 0 | 0 | 0 | 0 | 267260 | 7.16E+08 | 5.96E+08 | 6.86E+08 | 0 | 0 | 89086.67 | 6.66E+08 | D463H_D463N_(14) |
| 1401 | REV__E7EX40;REV__Q | REV__E7EX | 0 | 0 | 0 | 0 | 0 | 0 | 0 | 0 | 298600 | 318090 | 310090 | 291410 | 0 | 0 | 99533.33 | 306530 | D463H_D463N_(14) |
| 131 | W8JJB1;B4DS09;Q6PIS | BRD4;BRD3 | 0 | 0 | 0 | 0 | 0 | 0 | 683710 | 777230 | 0 | 0 | 0 | 0 | 0 | 0 | 486980 | 0 | D463H_only_(17) |
| 404 | A0A183YE00;A0A075B | A0A183YE( | 0 | 0 | 0 | 0 | 0 | 0 | 0 | 0 | 537500 | 0 | 0 | 0 | 0 | 0 | 179166.7 | 0 | D463H_only_(17) |
| 420 | Q53FA3;B2RCQ9;A0A1 | HSPA1L | 0 | 0 | 0 | 0 | 0 | 0 | 12846000 | 13348000 | 12599000 | 0 | 0 | 0 | 0 | 0 | 12931000 | 0 | D463H_only_(17) |
| 427 | H7C1C4;A0A223PQH6 | HCFC1 | 0 | 0 | 0 | 0 | 0 | 0 | 481130 | 438460 | 394790 | 0 | 0 | 0 | 0 | 0 | 438126.7 | 0 | D463H_only_(17) |
| 453 | A6H8W6;O43166;F5G\ | SIPA1L1 | 0 | 0 | 0 | 0 | 0 | 0 | 175840 | 0 | 174770 | 0 | 0 | 0 | 0 | 0 | 116870 | 0 | D463H_only_(17) |
| 567 | B2RTY4;H3BSU8;H3BM | MYO9A | 0 | 0 | 0 | 0 | 0 | 0 | 3484100 | 7157000 | 1550100 | 0 | 0 | 0 | 0 | 0 | 4063733 | 0 | D463H_only_(17) |
| 619 | B7ZA76;B4DIA9;Q8IX0 | UEVLD | 0 | 0 | 0 | 0 | 0 | 0 | 165640 | 159280 | 0 | 0 | 0 | 0 | 0 | 0 | 108306.7 | 0 | D463H_only_(17) |
| 694 | B7Z4B2;B7Z1V9;B7Z1H | NPEPPS | 0 | 0 | 0 | 0 | 0 | 0 | 247050 | 0 | 0 | 0 | 0 | 0 | 0 | 0 | 82350 | 0 | D463H_only_(17) |
| 735 | CON__O95678;O9567 | KRT75 | 0 | 0 | 0 | 0 | 0 | 0 | 569480 | 0 | 652550 | 0 | 0 | 0 | 0 | 0 | 407343.3 | 0 | D463H_only_(17) |
| 752 | CON__Q14525;Q1452 | KRT33B;KR | 0 | 0 | 0 | 0 | 0 | 0 | 417750 | 0 | 0 | 0 | 0 | 0 | 0 | 0 | 139250 | 0 | D463H_only_(17) |
| 860 | H3BLY5;H0YMM4;Q96/ | ISG20 | 0 | 0 | 0 | 0 | 0 | 0 | 760140 | 752540 | 0 | 0 | 0 | 0 | 0 | 0 | 504226.7 | 0 | D463H_only_(17) |
| 1019 | P17252-mutantDH | PRKCA | 0 | 0 | 0 | 0 | 0 | 0 | 99470000 | 1.06E+08 | 98262000 | 0 | 0 | 0 | 0 | 0 | 1.01E+08 | 0 | D463H_only_(17) |
| 1155 | Q08J23 | NSUN2 | 0 | 0 | 0 | 0 | 0 | 0 | 236140 | 252490 | 192280 | 0 | 0 | 0 | 0 | 0 | 226970 | 0 | D463H_only_(17) |
| 1228 | Q56G89 | Q56G89 | 0 | 0 | 0 | 0 | 0 | 0 | 590690 | 565270 | 685730 | 0 | 0 | 0 | 0 | 0 | 613896.7 | 0 | D463H_only_(17) |
| 1278 | Q7L3T8 | PARS2 | 0 | 0 | 0 | 0 | 0 | 0 | 268360 | 0 | 326460 | 0 | 0 | 0 | 0 | 0 | 198273.3 | 0 | D463H_only_(17) |
| 1342 | Q9BVA1 | TUBB2B | 0 | 0 | 0 | 0 | 0 | 0 | 535740 | 452320 | 432890 | 0 | 0 | 0 | 0 | 0 | 473650 | 0 | D463H_only_(17) |
| 1412 | X5D767 | ADA | 0 | 0 | 0 | 0 | 0 | 0 | 2962100 | 2837600 | 3508700 | 0 | 0 | 0 | 0 | 0 | 3102800 | 0 | D463H_only_(17) |
| 28 | A0A024R095;A0A024F | CLCC1 | 0 | 0 | 0 | 0 | 0 | 0 | 0 | 0 | 0 | 0 | 0 | 546240 | 0 | 0 | 0 | 182080 | D463N_only_(22) |
| 75 | A0A024R3P9;Q9H3P7 | ACBD3 | 0 | 0 | 0 | 0 | 0 | 0 | 0 | 0 | 0 | 0 | 0 | 529090 | 0 | 0 | 0 | 176363.3 | D463N_only_(22) |
| 163 | A0A024R9F5;P55040;I | GEM | 0 | 0 | 0 | 0 | 0 | 0 | 0 | 0 | 0 | 1078500 | 0 | 0 | 0 | 0 | 0 | 359500 | D463N_only_(22) |
| 226 | B4DKE6;Q5U0C2;Q53I | RFC3 | 0 | 0 | 0 | 0 | 0 | 0 | 0 | 0 | 0 | 0 | 418230 | 0 | 0 | 0 | 0 | 139410 | D463N_only_(22) |
| 265 | E5RHC7;E5RH59;E5RF | EIF3H;EIF3 | 0 | 0 | 0 | 0 | 0 | 0 | 0 | 0 | 0 | 346620 | 0 | 0 | 0 | 0 | 0 | 115540 | D463N_only_(22) |
| 332 | A0A0J9YW50;A0A0J9Y\ | A0A0J9YW5 | 0 | 0 | 0 | 0 | 0 | 0 | 0 | 0 | 0 | 115100 | 0 | 0 | 0 | 0 | 0 | 38366.67 | D463N_only_(22) |
| 409 | A0A1B0GUT5;Q8IYJ2 | C10orf67 | 0 | 0 | 0 | 0 | 0 | 0 | 0 | 0 | 0 | 0 | 0 | 763560 | 0 | 0 | 0 | 254520 | D463N_only_(22) |
| 502 | A8K8J9;H0YHL1;F8W1I | DCTN2 | 0 | 0 | 0 | 0 | 0 | 0 | 0 | 0 | 0 | 0 | 0 | 185730 | 0 | 0 | 0 | 61910 | D463N_only_(22) |
| 516 | B0AZS5;B4DM85;O001 | KIF2A | 0 | 0 | 0 | 0 | 0 | 0 | 0 | 0 | 0 | 0 | 664940 | 0 | 0 | 0 | 0 | 221646.7 | D463N_only_(22) |
| 747 | CON__P17690 | CON__P17( | 0 | 0 | 0 | 0 | 0 | 0 | 0 | 0 | 0 | 0 | 481090 | 0 | 0 | 0 | 0 | 160363.3 | D463N_only_(22) |
| 794 | E7DVW5;Q01469;I6L8 | FABP5 | 0 | 0 | 0 | 0 | 0 | 0 | 0 | 0 | 0 | 2715600 | 1908500 | 2828200 | 0 | 0 | 0 | 2484100 | D463N_only_(22) |
| 831 | F8VVR8;Q96LU7 | MYRFL | 0 | 0 | 0 | 0 | 0 | 0 | 0 | 0 | 0 | 0 | 0 | 111670 | 0 | 0 | 0 | 37223.33 | D463N_only_(22) |
| 848 | H7C2V8;H0Y765;H7BY | KMT2C | 0 | 0 | 0 | 0 | 0 | 0 | 0 | 0 | 0 | 0 | 4054300 | 5639500 | 0 | 0 | 0 | 3231267 | D463N_only_(22) |
| 869 | H7C0U1 | IWS1 | 0 | 0 | 0 | 0 | 0 | 0 | 0 | 0 | 0 | 495980 | 863450 | 636520 | 0 | 0 | 0 | 665316.7 | D463N_only_(22) |
| 988 | P05771 | PRKCB | 0 | 0 | 0 | 0 | 0 | 0 | 0 | 0 | 0 | 0 | 129780 | 0 | 0 | 0 | 0 | 43260 | D463N_only_(22) |
| 1056 | P31947 | SFN | 0 | 0 | 0 | 0 | 0 | 0 | 0 | 0 | 0 | 1612400 | 1524100 | 1474100 | 0 | 0 | 0 | 1536867 | D463N_only_(22) |
| 1096 | P53621 | COPA | 0 | 0 | 0 | 0 | 0 | 0 | 0 | 0 | 0 | 0 | 618360 | 507440 | 0 | 0 | 0 | 375266.7 | D463N_only_(22) |
| 1198 | Q15084 | PDIA6 | 0 | 0 | 0 | 0 | 0 | 0 | 0 | 0 | 0 | 231560 | 0 | 0 | 0 | 0 | 0 | 77186.67 | D463N_only_(22) |
| 1212 | Q6IBJ4;Q5U0N0;Q165 | SGCB | 0 | 0 | 0 | 0 | 0 | 0 | 0 | 0 | 0 | 5279400 | 0 | 0 | 0 | 0 | 0 | 1759800 | D463N_only_(22) |
| 1280 | Q7Z727 | PRKCA | 0 | 0 | 0 | 0 | 0 | 0 | 0 | 0 | 0 | 0 | 0 | 972560 | 0 | 0 | 0 | 324186.7 | D463N_only_(22) |
| 1305 | Q8NHU6 | TDRD7 | 0 | 0 | 0 | 0 | 0 | 0 | 0 | 0 | 0 | 234400 | 0 | 0 | 0 | 0 | 0 | 78133.33 | D463N_only_(22) |
| 1384 | Q9Y385;B4DUF8 | UBE2J1 | 0 | 0 | 0 | 0 | 0 | 0 | 0 | 0 | 0 | 0 | 0 | 160090 | 0 | 0 | 0 | 53363.33 | D463N_only_(22) |
| 15 | B4DSN3;G5EA31;A0A( | SEC24C | 0 | 0 | 0 | 898760 | 1077600 | 543420 | 0 | 946790 | 0 | 0 | 0 | 0 | 0 | 839926.7 | 315596.7 | 0 | WT_D463H_(22) |
| 127 | C9JKI3;Q7Z4F3;A0A02 | CAV1 | 0 | 0 | 0 | 620020 | 458900 | 499080 | 386480 | 386730 | 418220 | 0 | 0 | 0 | 0 | 526000 | 397143.3 | 0 | WT_D463H_(22) |
| 157 | B2ZWH1;B4DFD3;Q6N | CACYBP | 0 | 0 | 0 | 712570 | 0 | 0 | 432200 | 0 | 419500 | 0 | 0 | 0 | 0 | 237523.3 | 283900 | 0 | WT_D463H_(22) |
| 211 | F6S6P2;X6REW1;A0A0 | BAG6;BAT3 | 0 | 0 | 0 | 399700 | 270800 | 279160 | 274870 | 414150 | 325260 | 0 | 0 | 0 | 0 | 316553.3 | 338093.3 | 0 | WT_D463H_(22) |
| 279 | I3L3X7;A0A087X2I4;Q | PSMB6 | 0 | 0 | 0 | 1046200 | 1392600 | 1050900 | 0 | 1251800 | 0 | 0 | 0 | 0 | 0 | 1163233 | 417266.7 | 0 | WT_D463H_(22) |
| 392 | A0A140VK46;P28070; | PSMB4 | 0 | 0 | 0 | 0 | 303450 | 250970 | 0 | 0 | 299680 | 0 | 0 | 0 | 0 | 184806.7 | 99893.33 | 0 | WT_D463H_(22) |
| 432 | CON__P13646-1;A8K2 | KRT13 | 0 | 0 | 0 | 4750500 | 4038000 | 3499900 | 16214000 | 14043000 | 15610000 | 0 | 0 | 0 | 0 | 4096133 | 15289000 | 0 | WT_D463H_(22) |
| 499 | H3BM93;A8K897;H3B\ | NUP93 | 0 | 0 | 0 | 147300 | 162230 | 159900 | 165480 | 0 | 155600 | 0 | 0 | 0 | 0 | 156476.7 | 107026.7 | 0 | WT_D463H_(22) |
| 530 | B2R4M6;P06702 | S100A9 | 0 | 0 | 0 | 1256300 | 549480 | 0 | 1469800 | 1015000 | 0 | 0 | 0 | 0 | 0 | 601926.7 | 828266.7 | 0 | WT_D463H_(22) |
| 642 | B4DNG6;E9PHT9;V9H\ | ANXA5;HEL | 0 | 0 | 0 | 0 | 212310 | 0 | 301680 | 236680 | 298550 | 0 | 0 | 0 | 0 | 70770 | 278970 | 0 | WT_D463H_(22) |
| 648 | B4DPY2;H7C2N3 | B4DPY2;H7 | 0 | 0 | 0 | 3144200 | 12788000 | 0 | 0 | 737120 | 0 | 0 | 0 | 0 | 0 | 5310733 | 245706.7 | 0 | WT_D463H_(22) |
| 664 | E7EU94;E7ESX0;E7EN | PRPF31 | 0 | 0 | 0 | 0 | 0 | 71674 | 0 | 167160 | 132300 | 0 | 0 | 0 | 0 | 23891.33 | 99820 | 0 | WT_D463H_(22) |
| 679 | B4E2Z3;F5GZS6;J3KPF | SLC3A2 | 0 | 0 | 0 | 67152 | 71390 | 98519 | 99171 | 137230 | 99599 | 0 | 0 | 0 | 0 | 79020.33 | 112000 | 0 | WT_D463H_(22) |
| 727 | C9JL85;Q69YG1;P585 | MTPN;DKF | 0 | 0 | 0 | 0 | 1266800 | 0 | 0 | 0 | 709350 | 0 | 0 | 0 | 0 | 422266.7 | 236450 | 0 | WT_D463H_(22) |
| 837 | H7C5R5;F8WAE5;Q9B\ | EIF2A | 0 | 0 | 0 | 0 | 0 | 300800 | 0 | 188370 | 0 | 0 | 0 | 0 | 0 | 100266.7 | 62790 | 0 | WT_D463H_(22) |
| 862 | H3BQZ7;Q1KMD3 | HNRNPUL2 | 0 | 0 | 0 | 200700 | 359290 | 242100 | 224310 | 293740 | 277670 | 0 | 0 | 0 | 0 | 267363.3 | 265240 | 0 | WT_D463H_(22) |
| 1008 | P12035;CON__P1203 | KRT3 | 0 | 0 | 0 | 2292700 | 1413100 | 1876700 | 1689500 | 2068900 | 2088600 | 0 | 0 | 0 | 0 | 1860833 | 1949000 | 0 | WT_D463H_(22) |
| 1095 | Q5BKZ2;P52294;C9J3 | KPNA1 | 0 | 0 | 0 | 0 | 0 | 69901 | 111430 | 212320 | 290690 | 0 | 0 | 0 | 0 | 23300.33 | 204813.3 | 0 | WT_D463H_(22) |
| 1129 | Q59GN2;P62891 | RPL39P5;R | 0 | 0 | 0 | 1604100 | 0 | 0 | 2278200 | 2054900 | 2661500 | 0 | 0 | 0 | 0 | 534700 | 2331533 | 0 | WT_D463H_(22) |
| 1248 | Q5TDE9;Q5TDF0;Q9BS | C1orf57;NT | 0 | 0 | 0 | 0 | 0 | 383090 | 386400 | 0 | 417410 | 0 | 0 | 0 | 0 | 127696.7 | 267936.7 | 0 | WT_D463H_(22) |
| 1383 | Q9Y355;F8W696;A0A0 | APOA1 | 0 | 0 | 0 | 325490 | 306750 | 0 | 17344000 | 0 | 1.45e+09 | 0 | 0 | 0 | 0 | 210746.7 | 8672000 | 0 | WT_D463H_(22) |
| 1386 | Q9Y3C8 | UFC1 | 0 | 0 | 0 | 0 | 0 | 525740 | 0 | 948720 | 0 | 0 | 0 | 0 | 0 | 175246.7 | 316240 | 0 | WT_D463H_(22) |
| 69 | A0A024R333;Q6UXN9 | TMEM113;\ | 0 | 0 | 0 | 0 | 108730 | 0 | 0 | 0 | 0 | 0 | 158090 | 0 | 0 | 36243.33 | 0 | 52696.67 | WT_D463N_(20) |
| 181 | A0A024RAV0;P54687; | BCAT1 | 0 | 0 | 0 | 131000 | 167660 | 133140 | 0 | 0 | 0 | 127760 | 100030 | 118500 | 0 | 143933.3 | 0 | 115430 | WT_D463N_(20) |
| 207 | K4DIB8;K4DIA8;E9PS9 | SLC25A22 | 0 | 0 | 0 | 0 | 849820 | 0 | 0 | 0 | 0 | 0 | 648900 | 0 | 0 | 283273.3 | 0 | 216300 | WT_D463N_(20) |
| 213 | Q86YY1;B4DJ25;Q9NS | DKFZp434I | 0 | 0 | 0 | 0 | 422130 | 0 | 0 | 0 | 0 | 651790 | 644950 | 0 | 0 | 140710 | 0 | 432246.7 | WT_D463N_(20) |
| 360 | E5RIP4;A0A0S2Z5C8;A | SLC39A14 | 0 | 0 | 0 | 0 | 0 | 124060 | 0 | 0 | 0 | 0 | 78159 | 64806 | 0 | 41353.33 | 0 | 47655 | WT_D463N_(20) |
| 376 | D6R9X8;Q59F03;B4E0 | ITGA3 | 0 | 0 | 0 | 903720 | 0 | 286580 | 0 | 0 | 0 | 0 | 475880 | 751230 | 0 | 396766.7 | 0 | 409036.7 | WT_D463N_(20) |
| 416 | A0A1S5UZ07;Q9Y490 | TLN1 | 0 | 0 | 0 | 253070 | 0 | 256820 | 0 | 0 | 0 | 0 | 0 | 340160 | 0 | 169963.3 | 0 | 113386.7 | WT_D463N_(20) |
| 537 | Q68DQ4;B2R5T5;P106 | DKFZp779I | 0 | 0 | 0 | 0 | 179960 | 213040 | 0 | 0 | 0 | 56099 | 0 | 42952 | 0 | 131000 | 0 | 33017 | WT_D463N_(20) |
| 564 | Q5RKT7;B2RDW1;P62 | RPS27A;HE | 0 | 0 | 0 | 0 | 1191400 | 1261300 | 0 | 0 | 0 | 596600 | 3574300 | 3996000 | 0 | 817566.7 | 0 | 2722300 | WT_D463N_(20) |
| 614 | B4DE59;C9JTX4;C9J82 | B4DE59;C9 | 0 | 0 | 0 | 0 | 0 | 1056800 | 0 | 0 | 0 | 1256200 | 1590900 | 1622500 | 0 | 352266.7 | 0 | 1489867 | WT_D463N_(20) |
| 771 | D3DRG7;O14531;Q5T( | DPYSL4 | 0 | 0 | 0 | 0 | 724680 | 0 | 0 | 0 | 0 | 215520 | 225640 | 0 | 0 | 241560 | 0 | 147053.3 | WT_D463N_(20) |
| 836 | F8W930;Q9Y6M1;B4DI | IGF2BP2 | 0 | 0 | 0 | 0 | 0 | 262490 | 0 | 0 | 0 | 393320 | 635430 | 469810 | 0 | 87496.67 | 0 | 499520 | WT_D463N_(20) |
| 873 | H7C4H2;Q549N5;Q9Y | SRPRB | 0 | 0 | 0 | 0 | 411020 | 0 | 0 | 0 | 0 | 0 | 0 | 465290 | 0 | 137006.7 | 0 | 155096.7 | WT_D463N_(20) |
| 903 | K7ELJ8;K7ESP1;Q9961 | DNAJC7 | 0 | 0 | 0 | 112940 | 0 | 0 | 0 | 0 | 0 | 0 | 0 | 85361 | 0 | 37646.67 | 0 | 28453.67 | WT_D463N_(20) |
| 969 | O95831;A0A140VK04; | AIFM1 | 0 | 0 | 0 | 275570 | 0 | 0 | 0 | 0 | 0 | 291220 | 0 | 0 | 0 | 91856.67 | 0 | 97073.33 | WT_D463N_(20) |
| 977 | P04259 | KRT6B | 0 | 0 | 0 | 0 | 467410 | 0 | 0 | 0 | 0 | 5460700 | 5983500 | 2765600 | 0 | 155803.3 | 0 | 4736600 | WT_D463N_(20) |
| 1050 | P29508;Q5K684;Q5K6 | SERPINB3 | 0 | 0 | 0 | 1215000 | 0 | 0 | 0 | 0 | 0 | 0 | 1274000 | 0 | 0 | 405000 | 0 | 424666.7 | WT_D463N_(20) |
| 1079 | P47929 | LGALS7 | 0 | 0 | 0 | 523420 | 806470 | 904530 | 0 | 0 | 0 | 5167600 | 2118600 | 1716400 | 0 | 744806.7 | 0 | 3000867 | WT_D463N_(20) |
| 1112 | P61964;V9GYQ5;V9GZ | WDR5 | 0 | 0 | 0 | 131290 | 0 | 0 | 0 | 0 | 0 | 187890 | 0 | 0 | 0 | 43763.33 | 0 | 62630 | WT_D463N_(20) |
| 1244 | Q5T749 | KPRP | 0 | 0 | 0 | 609900 | 422410 | 0 | 0 | 0 | 0 | 576730 | 336360 | 643770 | 0 | 344103.3 | 0 | 518953.3 | WT_D463N_(20) |
| 175 | B7Z881;Q59HA3;E7EW | IQGAP2 | 0 | 0 | 0 | 0 | 321800 | 167930 | 0 | 0 | 0 | 0 | 0 | 0 | 0 | 163243.3 | 0 | 0 | WT_only_(12) |
| 403 | H0YBR5;A0A161SXE1; | EIF3E | 0 | 0 | 0 | 98609 | 0 | 0 | 0 | 0 | 0 | 0 | 0 | 0 | 0 | 32869.67 | 0 | 0 | WT_only_(12) |
| 471 | G3V1S1;B3KTH1;B3KR | ERCC3 | 0 | 0 | 0 | 0 | 0 | 56822 | 0 | 0 | 0 | 0 | 0 | 0 | 0 | 18940.67 | 0 | 0 | WT_only_(12) |
| 572 | B3KMK7;Q9Y2A7 | NCKAP1 | 0 | 0 | 0 | 0 | 286870 | 617470 | 0 | 0 | 0 | 0 | 0 | 0 | 0 | 301446.7 | 0 | 0 | WT_only_(12) |
| 710 | Q9BUD9;E9PG46;D6W | AAK1;BMP2 | 0 | 0 | 0 | 0 | 67018 | 55651 | 0 | 0 | 0 | 0 | 0 | 0 | 0 | 40889.67 | 0 | 0 | WT_only_(12) |
| 721 | C9J8B8;Q969S8;Q59FI | HDAC10 | 0 | 0 | 0 | 0 | 125060 | 544130 | 0 | 0 | 0 | 0 | 0 | 0 | 0 | 223063.3 | 0 | 0 | WT_only_(12) |
| 893 | J3QLI9;Q7Z5A3;P6231 | SNRPD1 | 0 | 0 | 0 | 0 | 0 | 460540 | 0 | 0 | 0 | 0 | 0 | 0 | 0 | 153513.3 | 0 | 0 | WT_only_(12) |
| 999 | P0C7V6 | SUPT20HL | 0 | 0 | 0 | 0 | 0 | 157470 | 0 | 0 | 0 | 0 | 0 | 0 | 0 | 52490 | 0 | 0 | WT_only_(12) |
| 1197 | Q15061 | WDR43 | 0 | 0 | 0 | 0 | 6450500 | 0 | 0 | 0 | 0 | 0 | 0 | 0 | 0 | 2150167 | 0 | 0 | WT_only_(12) |
| 1235 | Q5CZA6;P53804;H7C( | DKFZp686I | 0 | 0 | 0 | 0 | 1719800 | 0 | 0 | 0 | 0 | 0 | 0 | 0 | 0 | 573266.7 | 0 | 0 | WT_only_(12) |
| 1243 | Q5T440 | IBA57 | 0 | 0 | 0 | 0 | 0 | 443710 | 0 | 0 | 0 | 0 | 0 | 0 | 0 | 147903.3 | 0 | 0 | WT_only_(12) |
| 1411 | W8QEH3 | LMNA | 0 | 0 | 0 | 0 | 720800 | 513600 | 0 | 0 | 0 | 0 | 0 | 0 | 0 | 411466.7 | 0 | 0 | WT_only_(12) |

**Table S13. Exclusive PKCα mutant D463H interactors.** Proteins exclusively co-immunoprecipitated with MYC-tagged D463H-PKCα. Only proteins identified in a minimum of 2 out of 3 D463H- PKCα replicate precipitates, which did not co-precipitate in any control, WT or D463N sample, are presented.

| ID | Name | MW (kDa) |
|---|---|---|
| **HSPA1L** | Heat Shock Protein Family A (Hsp70) Member 1 Like | 70 |
| **NSUN2** | NOP2/Sun RNA Methyltransferase 2 | 86 |
| **HCFC1** | Host Cell Factor | 208 |
| **TUBB2B** | Tubulin Beta 2B Class IIb | 50 |
| **ADA** | Adenosine Deaminase | 41 |
| **MYO9A** | Myosin IXA | 293 |
| **ISG20** | Interferon Stimulated Exonuclease Gene 20 | 20 |
| **UEVLD** | UEV And Lactate/Malate Dehydrogenase Domains | 52 |
| **BRD4:BRD3** | Bromodomain Containing 4: Bromodomain Containing 3 | 152 |
| **SIPA1L1** | Signal Induced Proliferation Associated 1 Like 1 | 200 |
| **PARS2** | Prolyl-TRNA Synthetase 2, Mitochondrial | 53 |
